# Corticothalamic modelling of sleep neurophysiology with applications to mobile EEG

**DOI:** 10.1101/2024.02.28.582655

**Authors:** Taha Morshedzadeh, Kevin Kadak, Sorenza P. Bastiaens, M. Parsa Oveisi, Davide Momi, Zheng Wang, Shreyas Harita, Maurice Abou Jaude, Christopher A. Aimone, Steve Mann, Sean L. Hill, John D. Griffiths

## Abstract

Recent developments in mathematical modelling of EEG enable the tracking of otherwise-inaccessible neurophysiological parameters throughout sleep. Likewise, advancements in wearable electronics have enabled easy & affordable collection of sleep EEG at home. The convergence of these two advances, namely neurophysiological modelling for mobile sleep EEG, can boost preclinical and clinical assessments of sleep. However, this subject area has received limited attention in existing literature. To address this, we used an established model of the corticothalamic system to analyze EEG power spectra from 5 datasets, spanning from research-grade systems to at-home mobile EEG. In the present work, we compare the convergent and divergent features of the data and the estimated physiological model parameters. While data quality and characteristics differ considerably, key patterns consistent with previous theoretical and empirical work are observed. During the transition from lighter to deeper NREM, i) exponent of the aperiodic (1*/f*) spectral component is increased, ii) bottom-up thalamocortical drive is reduced, iii) corticocortical connection strengths are increased. This effect is observed in healthy subjects but is interestingly absent when taking SSRI antidepressants, suggesting possible effects of ascending neuromodulation on corticothalamic oscillations. We further show a month-long increase in REM% in one mobile EEG subject, associated with boosted high-frequency activity in spectra and higher thalamothalamic gains in the model, pointing to possible changes of thalamic inhibition in REM parasomnias. Our results provide a proof-of-principle for the utility and feasibility of this physiological modelling-based approach to analyzing mobile EEG data, providing a mechanistic measure of brain physiology during sleep.

**Statement of significance:** We employ a physiological model of the corticothalamic circuitry to model the EEG power spectra in sleep. We fit this model to 5 EEG datasets, and demonstrate that while mobile and non-mobile EEG recordings differ in their characteristics and quality, they can both robustly represent the changes along sleep stages using the aperiodic (1*/f*) component. We observe an increased corticocortical connection strength and decreased corticothalamic connection strength as the subject goes into deeper stages of NREM sleep; an effect that is, importantly, not observed in subjects taking SSRIs. This work provides a proof-of-concept for using mathematical modelling, working well for large mobile and non-mobile datasets providing valuable insight into the mechanisms generating sleep EEG.

## 1 Introduction

### Sleep neurophysiology and EEG

Sleep is a vital and near-universal physiological function, manifested in most of the animal kingdom in a regular circadian pattern [1, 2]. It is far more than just a state of rest and reduced energy expenditure. States of sleep serve as a period during which the brain undergoes significant changes, including metabolic homeostasis and recovery [3, 4], synaptic regulation, and memory consolidation [1, 5]. Disturbances in sleep rhythms can also increase susceptibility to various types of psychiatric and neurological conditions such as mood disorders, epilepsy, and dementia [6–9]. Despite these associations, sleep disorders remain highly underdiagnosed clinically, or misdiagnosed as other neurological ailments [10].

The physiological state of the brain moves through a complex trajectory of dynamical regimes during a night’s sleep. These changes evolve on the timescale of tens of minutes, and their electrical footprints are reflected in (and are indeed defined by) electroencephalography (EEG) recordings. Polysomnography (PSG), is one of the most widely-used methods for evaluating sleep in the clinical setting. It involves the concurrent monitoring of EEG, electromyography (EMG), electrooculography (EOG), movement, and respiration. Sleep stages are defined in terms of the properties of EEG time series data over standard (30s) windows, and the time series of the stages for the successive windows forms the hypnogram. This data is typically evaluated over a single channel, following the –mostly correct– assumption that brain activity changes similarly across all EEG channels during sleep [11]. Each of these stages has characteristic phenomenological definitions defined by the common sleep staging standards [12, 13]. Although it has strong diagnostic and prognostic utility [14, 7, 15–17], classical sleep staging is highly constrained as it is limited to only 5 values (the 5 stages W, N1, N2, N3, REM) to capture the vast continuum of brain states in sleep. This problem is further exacerbated by the highly subjective interpretation of different stages by human scorers, which has led to considerable inter-expert variability [18, 19]. Therefore, it is crucial to augment this information with more detailed quantitative approaches for evaluating brain activity trajectories in sleep.

Power spectral estimation is one of the fundamental methods for studying the characteristics of a time series signal across different frequencies. Studying the EEG power spectral density (PSD) in the same 30-second windows used for sleep staging can provide us with a more high-dimensional evaluation of brain states over these intervals. EEG PSDs can be reliably described in terms of two main components: i) A background 1*/f*^*n*^ trend, understood to be non-oscillatory or ‘aperiodic’, and defined by its exponent and offset, and ii) An oscillatory component which is highly periodic, featuring well-defined attributes such as frequency, amplitude, and bandwidth. The aperiodic component is an intrinsic feature of many natural processes, and is believed in the neuroscientific context to reflect variable excitatory/inhibitory balance [20]. It has also been linked to cognitive decline in ageing [21], cognitive speed [22], and movement [23]. Periodic activity is traditionally examined in the frequency bands delta (0.5-4 Hz), theta (4-7.5 Hz), alpha (7.5-12 Hz), beta (16-30 Hz), and gamma (>30 Hz). During wakefulness, the brain exhibits high-frequency low-amplitude activity, and as the subject transitions to NREM sleep, the activity transitions into a low-frequency high-amplitude pattern.[24, 25]. The transition from lighter to deeper NREM sleep is associated with increased slope of the EEG power spectrum, along with increases in the amplitude of the delta band and a decrease in the amplitude the alpha band [26]. Wakefulness also is signified by a presence of alpha and gamma peaks, and REM (rapid eye movement) sleep is correlated with increases in gamma and theta but not alpha peaks [27].

The principal brain structure that drives brain state changes during sleep, including our measurement of them with EEG, is the corticothalamic system [28–30]. Different stages of sleep have been linked to changes in corticothalamic activity [29, 31–34] and to changes in the periodic and aperiodic components of the EEG signals over those changes [35, 36, 34, 37]. For instance, the transition from wakefulness to N1 sleep is also characterized by an increase in the slope of the 1*/f* component and the low-frequency band powers [35, 36], which is itself observed to be associated with corticothalamic communication [30].

Sleep stage N3, also known as slow-wave sleep (SWS), is understood as the deepest stage of NREM sleep, showing strongly synchronized cortical activity in the infra-slow (<1 Hz) and delta (1-4 Hz) frequency bands. This synchronized cortical activity has been shown to be driven locally through corticocortical connections, and with reduced thalamocortical input [31, 38–40]. Interestingly, 1 Hz transcranial magnetic stimulation (TMS) in the cortex can effectively entrain this 1 Hz cortical oscillation around the stimulation site [41], indicating that cortical activation is the primary source driving this oscillation. Synaptic homeostasis and long-term potentiation (LTP) have also been found to occur strongly in SWS [5, 33], and brain stimulation at this stage can trigger memory replays and improve memory recall [42].

### Mathematical modelling of sleep-wake dynamics

This deep foundation of experimental knowledge in neuroscience across multiple species, spatial scales, and observable phenomena, provides a strong motivation for the development and use of mathematical models that explain sleep EEG in terms of their underlying neurophysiological processes across the units of the corticothalamic circuitry. One of the most widely used and extensively studied models of this kind to date was introduced by Robinson et al. [30], which describes, at the mesoscopic spatial scale, a four-node corticothalamic network containing the thalamic relay, thalamic reticular, cortical excitatory, and cortical inhibitory neural populations. With this structure, the Robinson model has proved highly capable of replicating measured EEG time series and power spectra [30, 43–45], with applications including evoked potentials [46, 47], alpha rhythms [48], and sleep & arousal [49, 50], to name only a few. In a 2015 paper, Abeysuriya et al. demonstrate the use of this model to study the trajectories of physiological brain states expressed through the EEG, across a night of sleep [51]. By fitting the model-generated power spectra to those observed in empirical EEG, circuit mechanisms such as corticocortical, corticothalamic, and intrathalamic connection strengths can be estimated from 30-second windows rolling throughout the night, and their changes compared against separately-scored PSG classifications. In this way, mathematical modelling of corticothalamic system dynamics can be used to enrich the observations made via classical sleep stages and conventional power spectral analysis.

### Emerging mobile neurotechnologies for sleep EEG measurements

One of the most significant technical developments in the field of EEG over the past decade has been a suite of hardware, software, and commercial innovations leading to the widespread availability of low-cost (“consumer-grade”) wireless mobile EEG devices. The lower price tag, smaller footprint, use of flexible components such as conductive rubber and conductive fabric, and more streamlined setup of these systems hold great promise for scientists and clinicians needing to access larger samples of subjects and over many more nights than is possible with traditional in-lab sleep EEG assessments. Two of the most established mobile sleep EEG headsets on the market today are Muse S by InteraXon [52] and Dreem by Beacon Biosignals [53]. Although, these products face stiff competition from other startups that with smaller but increasing market share, such as Cerebra [54], URGOnight [55], IDUN [56], and Elemind [57], along with major consumer electronics companies such as LG Electronics (sleepwave.ai) that are looking to enter the mobile sleep EEG market.

This approach can enable an easier and more affordable overnight recording of sleep EEG at home or in the research lab. The easier setup and reduced cost can readily enable the researchers to make recordings over more repeated nights and for a larger population.

### Characterizing trajectories of activity in healthy vs. unhealthy sleep

The mathematical models of EEG activity enable us to reconstruct an embedding space underlying the changes in EEG activity observed in sleep. Fitting these models to repeated recordings from the a larger sample size of participants enable us to catalogue a rich set of ranges and the trajectories of the physiological parameters from the model in various nights of sleep. Not only can applying such mathematical models to repeated recordings from a larger sample size of participants help us characterize the ranges of normative parameters correlated with good restorative sleep, but the repeated recordings can also help us detect the ranges associated with sporadic changes in sleep quality or potential parasomnias that require continuous monitoring [58, 59].

Certain sleep EEG patterns are correlated with mood, anxiety, and other mental health factors, but this area remains understudied due to the logistical challenges of the repeated recording of sleep EEG over extended periods, especially from subclinical, at-risk, or asymptomatic populations who are at home rather than in controlled, hospitalized settings [60–62]. Mobile EEG systems are key in bridging this gap, since they make continuous and long-term monitoring of sleep EEG outside of the clinical environment feasible.

Additionally, there is significant night-by-night variability in sleep within the same individual. Collecting extensive nightly data from the same person allows the identification of consistent, robust patterns unique to that individual, and it reduces the effects of these stochastic fluctuations. Dreaming is an example of a sparse sleep event which varies night-by-night, is associated with many determinants of mental and physical health [63, 64], and its actuation is strongly affected by the level of comfort in sleep. These factors make it a prime example of a topic that is best investigated using large-sample-size longitudinal mEEG recordings, as evidenced by ongoing data collection projects such as [65] Brain states undergo semi-regular trajectories and cycles of changes through a night of sleep—which we describe at a phenotypical level as the hypnogram of the sleep stages. These variations are reflected by the changes in aperiodic and periodic components of the EEG activity in the power spectral domain. Therefore, we can attain a trajectory of the parameters underlying those brain states suggested by the corticothalamic model. Beyond mapping these parameter trajectories to various health or disease states, various interventions and treatments can alter the ranges and trajectories of these parameters in a unique way, which could be captured via the parameter trajectories describing underlying physiological state transitions of the brain.

In summary, by fitting many such sleep recordings to the mathematical model, we can characterize the embeddings and their transitions associated with *healthy* sleep, and detect canonical patterns of activity associated with this state. Moreover, we can examine parameters derived from fitting the model to *unhealthy* sleep EEG to understand how these key patterns deviate and where disruptions occur. And lastly, we can observe again how various types of *interventions* can change brain activity.

### Personalized medicine informed by physiological modelling of mEEG data

In the recent years, there has been a welcome shift in the computational neuroscience towards implementing the mathematical models of brain activity to *simulate* an individual’s brain activity in health and sleep. This has especially been explored in brain stimulation research where customized simulations of each person’s brain, informed by its connectomics, are used to predict the effects of the stimulation that is to be delivered. Lang et al. [66] provide a thorough review of such approaches in Neurosurgery. For instance, a “Virtual Epilepsy Patient” can be simulated to help detect the epileptogenic zone and devise various surgical and therapeutic interventions [67, 68].

The benefits of such modelling approaches is not just limited to the clinical implementation by the bedside. Rather, it can even be used to assist with the development of new therapeutic choices. An example of such work is demonstrated by Haas et al. [69], where *in-silico* simulated experiments using biophysical models of the human cortex correctly predicted the inefficacy of a certain new drug in trial even better than the animal models the drugs where tested on.

Utilizing the data from each individual, we can build a *personalized* simulation of their brain in sleep, which has a customized range and set of properties associated with their sleep. This can not only assist with the diagnostic process, but can also enlighten us on the underlying processes giving rise to these drops in sleep quality, and also help design new treatments and monitor & predict the prognostics of the treatment response.

### Present work

The recent advances cited above in our fundamental understanding of sleep neurophysiology, our ability to formulate and model it mathematically, and in the emergence of new technologies promising to radically up-scale the accessibility of EEG-based sleep monitoring, prompt a series of important research questions at the intersection of these topics. Previous work on personalized medicine through mesoscopic modelling of the brain has been limited to data that is collected in lab and in clinical departments–and we aimed to study whether we can utilize mobile EEG in the same way to develop personalized models of brain activity in health & disease.

This was the focus of the present study. Selecting several widely used open-access research- and consumer-grade sleep EEG datasets, we used power spectral analysis to first evaluate the changes to periodic and aperiodic spectrum components across sleep stages during the night of sleep, assessing the performance of different EEG systems in capturing variations in physiological brain states. We then fit the Robinson corticothalamic model to these EEG power spectra, with a view to studying mechanisms underlying these physiological states over sleep stages, and evaluating their correlations to the depth of NREM sleep. Lastly, we used health data from one of the analyzed cohorts to investigate the correlations of model-estimated neurophysiological parameters with specific mental & physical health biomarkers.

## 2 Methods

### 2.1 EEG Datasets

We used EEG data from multiple sources, described in the following. All datasets were acquired according to the ethics board regulation at the hosting institutions. They were accessed and used in accordance with their relevant licences and data-sharing agreements. The left frontal, central, or temporoparietal channels were used in each dataset, specifically F3 or adjacent 10/20 system locations, subject to availability and data quality.

#### 2.1.1 Sleep European Data Format - Extended (Sleep-EDFX)

We used 197 recordings from 185 subjects (97 female / 78 male, mean age 54.7), which were recorded in the time span of 1987-1991 and 1994 using portable Walkman-style cassettes at home [70, 71]. The accessed data had been digitized from the analog signal at the sampling rate of 100 Hz. In this dataset, 153 of the subjects had no previous health conditions and 44 were generally healthy but had trouble sleeping. We accessed the dataset through PhysioNet [72], acquiring the version last updated in 2018. We selected the data from the Fpz-Cz electrode channel for this work. The sleep stages were originally marked according to the Rechtschaffen & Kales (R&K) method [13], and were transformed into the AASM standard for further use in this project. In this paper, we will refer to this dataset as EDF-X for brevity.

#### 2.1.2 Dreem Open Datasets (DOD)

This dataset includes 80 PSGs collected using a research-grade PSG setup from 80 subjects (54 male / 26 female, mean age 42.39). The dataset was curated by Dreem, a manufacturer of sleep EEG headsets, to benchmark automatic sleep staging methods [73, 74]. The data was sampled at 250 Hz and scored by sleep professionals based on the 2007 AASM manual. This dataset is comprised of two sections: 25 healthy subjects recorded in Bretigny-Sur-Orge, France, over 12 mastoid-referenced EEG channels; and 55 at Redwood City, CA, USA, from subjects with Obstructive Sleep Apnea (OSA), with 8 mastoid-referenced EEG channels. We used the Fp2-O2 channel data from this dataset for the current project.

#### 2.1.3 Nap-EEG Dataset

This dataset was acquired from a healthy cohort of 22 individuals (16 male / 6 female; mean age 25.5 ± 7.03). The data was recorded between 10:00 and 17:00 over 1 to 2 consecutive days at the City College of New York, totalling 41 recordings. Subjects took a high- or low-load cognitive task and then took a nap for 30 minutes, where their EEG would be recorded over 64 channels of EEG and 2 channels of EOG, sampled at 1.5 KHz [75]. The data includes sleep stages and 2528 sleep spindles manually annotated, according to the AASM system. The dataset was accessed through the Open Science Framework (OSF) [76]. The PO8 channel data was chosen for the purpose of this project.

#### 2.1.4 Wisconsin Sleep Cohort (WSC) Dataset

The WSC dataset is recorded from a large cohort of state employees in Wisconsin, United States. We requested and accessed the standardized dataset through the National Sleep Research Resource (NSRR) [77, 78]. This set includes 2570 recordings from 1123 subjects. This is a longitudinal data set in which the same subjects came to the sleep lab every 4 to 5 years for a PSG recording. Each subject has 1 to 5 EEG recordings through the years, each approximately 4.5 years apart on average (mean 4.54 ± 1.50 years) between 2000 and 2015. The percentage of subjects with 1, 2, 3, 4, and 5 recordings was respectively 32.5%, 17.1%, 39.6%, 10.6%, and 0.2%. The subjects were 37 to 85 years old (mean age 59.82 ± 8.49, 1385 male / 1185 female). The 6 recording EEG electrodes were referenced to the ipsilateral mastoid electrodes and sampled at 100 Hz for the data from 2000 - 2009, and at 200 Hz from 2009 to 2015. This dataset also includes a large variety of mental and physical health information, such as the Zung Depression Scale, anxiety scales, caffeine consumption, number of recent nights with insomnia, blood pressure disorders, current medications, etc. We used the 556 first recordings in this dataset from 248 subjects, 39 to 81 years of age (mean age 60.29 ± 8.62, 137 male / 111 female). The C3 channel data was chosen for the purpose of this project.

#### 2.1.5 Muse’s Sleep Dataset (MSD)

Muse S is a wireless sleep EEG headband manufactured by InteraXon (Toronto, ON, Canada). The data is sampled at 256 Hz, recorded from dry electrodes TP9, TP10, Fp1 and Fp2, and referenced at the FpZ electrode. We used 10 recordings made between 2020 and 2022, provided by InteraXon, that were selected from Muse’s Sleep Dataset (MSD). MSD is an internal dataset of overnight at-home sleep recordings collected with the Muse S EEG headset [52]. This dataset was collected in accordance with the privacy policy that users agree to when using the Muse headband and ensures their informed consent concerning the use of EEG data for scientific research purposes. The subjects were 26 to 68 years old (mean age 38.70 ± 13.07, 7 male / 3 female) at the time of recordings. Sleep stages for these recordings were produced by Muse’s proprietary automated sleep staging algorithm [52]. Due to the increased presence of artifacts in the data recorded from dry EEG electrodes [79], for each subject, we marked the 30s epochs with a standard deviation larger than that of the whole recording, thus dropping an approximate 7.48% of all epochs from power spectral analysis & model fitting across the entire dataset. In this work, we use the EEG data in the TP7 channel from the MSD dataset, as it uses a frontal reference, thus quantifying the differential trace between the temporoparietal and frontal electrodes.

We also obtained a second set of 78 EEG recordings from 2 Muse S users, to use as a case study examining the suitability of such repeated nightly EEG recordings for monitoring sleep health and brain activity. These 2 users recorded their sleep at least every other night for a total period of 30-60 days. User 1 has 32 recordings and user 2 has 46 recordings. All processing steps were done similar to the MSD data described above. These 78 recordings were not used for the general group-level analyses, as they include a mix of normal and abnormal sleep parameters across various nights.

### 2.2 EEG & Hypnogram Data Analysis

#### 2.2.1 Pre-processing

The data were organized and processed using MNE-Python [80]. The power spectral density from the data was calculated in 30s segments using Welch’s method [81] in 4s Hamming windows with a 1s overlap. Choice of window lengths in EEG signal processing should be optimized for the analysis objectives in question [82]. In the present case, this choice of window sizes was made to balance the sharpness of the peak frequencies –due to noise-driven changes in the power of those bands– with a physiologically-plausible level of specificity in key rhythms such as alpha (7.5-12 Hz). And the segment length here was chosen as it is the segment length over which the sleep stages are labelled. The sleep staging system used here is the 2007 standard, issued by the American Academy of Sleep Medicine (AASM) [12]. The epochs with stages marked as *unknown* were omitted for all datasets.

#### 2.2.2 Aperiodic component estimation using FOOOF

We used the Python library `FOOOF v1.0.0` to separate the periodic and aperiodic components of the empirical and fitted power spectra [83]. The algorithm fit a Gaussian power spectrum corresponding to the aperiodic component to each EEG PSD without a knee, in the range of 0-45 Hz, with bins of the size 0.25 Hz. The Gaussian spectrum was then deducted from the EEG power spectrum to separate the periodic (oscillatory) components. This process was implemented iteratively and optimized to get maximum 4 oscillatory peaks, each between 1 to 4 Hz in bandwidth, and with at least 1 V^2^ / Hz amplitude. We used the extracted exponent (slope) & offset of the fitted aperiodic (1*/f* -like) component and the frequency & power of the periodic components to study the phenomenological properties of the power spectra.

#### 2.2.3 Calculating sleep metrics and statistics using YASA

We used the Python library `YASA v0.6.3` to estimate some common sleep metrics based on the presented hypnograms [84]. This library calculated values for sleep architecture and quality metrics using the subjects’ hypnograms according to the AASM guidelines [12]. Table 1 includes these evaluated metrics.

**Table 1:** Abbreviations and descriptions of sleep the hypnogram-based sleep metrics calculated using YASA.

| Metric | Description | Unit |
| --- | --- | --- |
| TIB | Total duration of the hypnogram. | min |
| SPT | Total duration from first to last period of sleep. | min |
| WASO | Duration of wake periods between the first and last periods of sleep | min |
| TST | Total duration of N1 + N2 + N3 + REM sleep between the first and last periods of sleep. | min |
| SE | $(TST / TIB) \times 100$ | % |
| SME | $(TST / SPT) \times 100$ | % |
| SOL | Latency to first epoch of sleep | min |
| <Stage> Latency | Latency to the first instance of the sleep stage <Stage> | min |

**Table 2:**
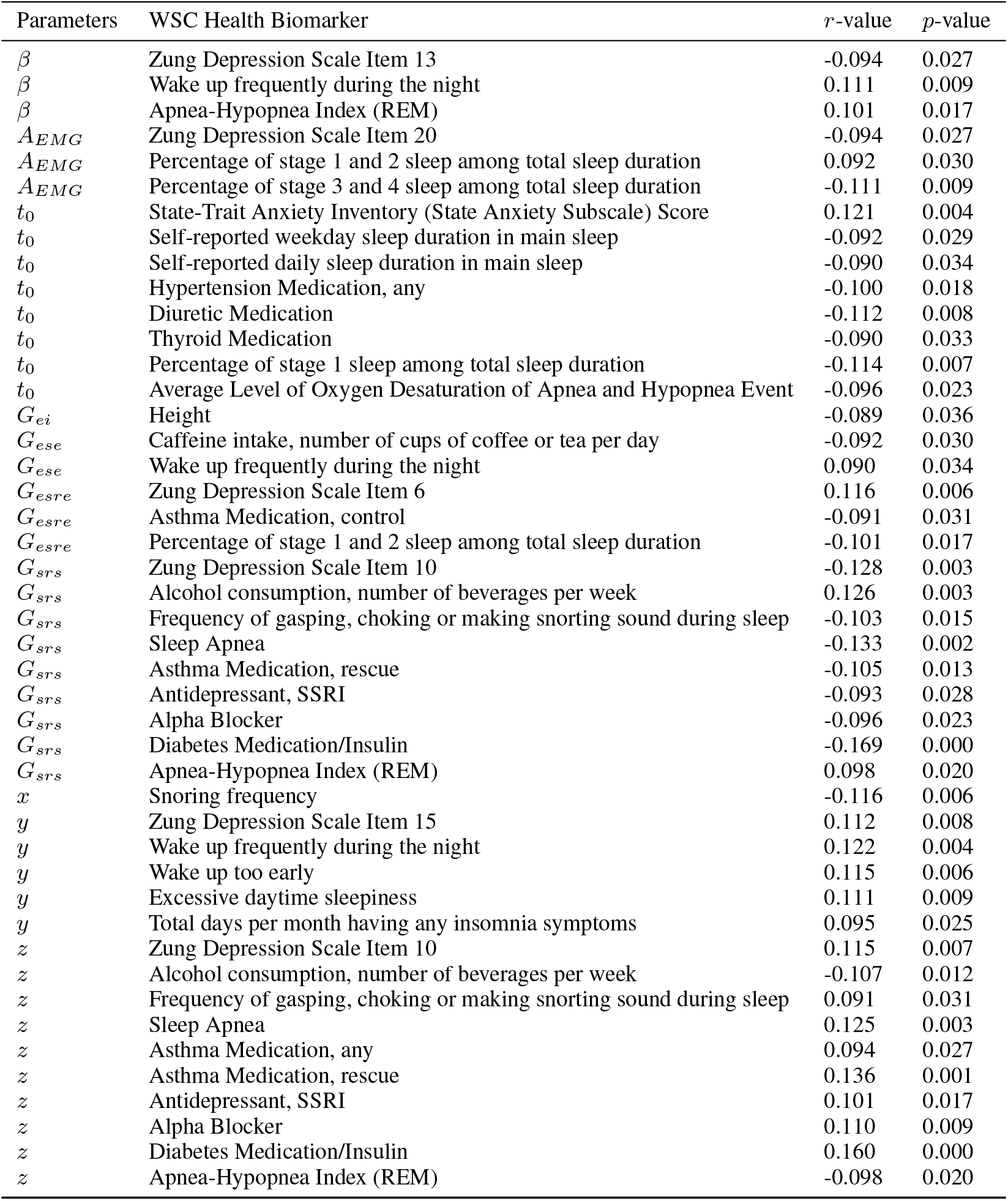
Significant correlations between biomarkers and average parameters per night in WSC.

### 2.3 Neurophysiological Model of Thalamocortical Activity

This work used a *neural field model* of thalamocortical dynamics to simulate plausible activity observed in the EEG data [30, 45]. In this physiological wave equation model, we model activity across these units of the thalamocortical circuitry, generated by: cortical excitatory (pyramidal) neurons (*e*), cortical inhibitory interneurons (*i*), thalamic reticular nucleus (*r*) and thalamic relay nuclei (*s*).

In each of these populations, a mean firing rate (i.e., pulse density) of each population denoted as *Q*_*a*_ is calculated based on the mean somatic voltage (*V*_*a*_). Henceforth in this document, *a & b* will represent any of the modelled populations, *a, b* ∈ {*e, i, r, s*}.

With *θ*_*a*_ as the mean firing threshold and 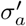 as the standard deviation of the somatic voltage for the population *a*, we can calculate *Q*_*a*_ as:

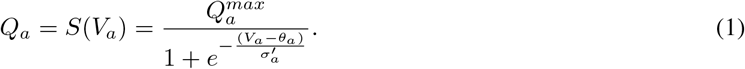

Using *Q*, the number of outgoing axonal spikes from the population (*ϕ*_*a*_) can be determined via this equation:

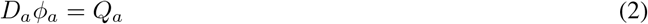

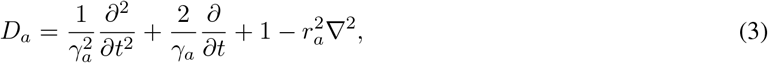

where *D*_*a*_ is a nonlinear term that dampens the incoming spike rate into a *field* equation (*ϕ*), and the temporal damping rate is *γ*_*a*_ = *υ*_*a*_*/r*_*a*_. For each population *a, υ*_*a*_ is the axonal conduction velocity, which is approximated to 10 *m/s* for myelinated axons that form the thalamocortical projections. *r*_*a*_ is the total range of axons of type *a*. This model assumes that long-range connections are myelinated and hence have a higher *υ*_*a*_. Shorter-range connections are not myelinated and will only have negligible values, of *υ*, rendering their effects on cortical activity insignificant. Among the thalamocortical connections, only the thalamic relay-excitatory cortical (*r* ↔ *e*) connection has a non-negligible distance, and as a result, we can take *r*_*e*_ as the only significant *r* value affecting the propagation and assume the other *r* values to be 1.

In the subcortical units (*r* and *s*), we also assume spatial uniformity such that ∇^2^ = 0. Given the large value of *γ*_*a*_ for the thalamocortical connection, the damper term (*D*_*a*_) converges to 1. As such, we can approximate that in the physiological states, *Q*_*a*_ = *ϕ*_*a*_.

To account for the delays introduced by the synapses, we introduce 1*/β* and 1*/α* which are respectively the *rise* and *decay* time constants of the postsynaptic soma activation, in the response of a population to a spike. We can write the dendritic impulse response function as:

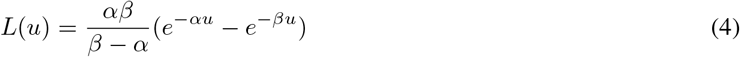

So, if *α* ≠ *β*, the dendritic activation function can be written as:

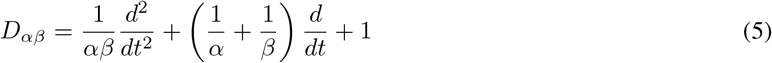

The Fourier transform of *L*(*u*) yields a function in which the dendritic impulse response acts as a low-pass filter with the cut-off frequency at *α*, and exhibiting a more steep attenuation at *β* Hz:

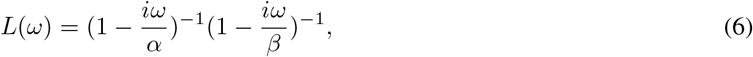

where *ω* = 2*πf* is the angular frequency and *f* is the frequency in Hz.

We take *V*_*a*_(*r, t*) to be the electrical field (voltage) in population *a*, influenced by: 1) *ϕ*_*ab*_ which is the incoming activation by the presynaptic populations from population *b*, 2) *N*_*ab*_ the mean number of synapses between *a* and *b*, and 3) *s*_*ab*_ the strength of each synapse between these two populations (the time-integrated response strength in the soma for each incoming spike):

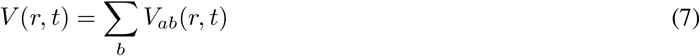

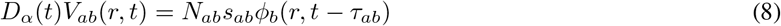

We further define *ν*_*ab*_ = *N*_*ab*_*s*_*ab*_ as the strength of all incoming synapses. The value of *s* (and hence *ν*) are considered positive for excitatory neurons and negative for inhibitory neurons. In this work, we assume *random connectivity* in the excitatory and inhibitory populations in the cortex, which means that *N*_*ia*_ = *N*_*ea*_ for any population *a*. Therefore, we simplify the *ν* values as follows: *ν*_*ee*_ = *ν*_*ie*_, *ν*_*ei*_ = *ν*_*ii*_, and *ν*_*es*_ = *ν*_*is*_. Hence, we are left with the 8 independent *ν* values: *ν*_*ee*_, *ν*_*ei*_, *ν*_*es*_, *ν*_*se*_, *ν*_*sr*_, *ν*_*rs*_, *ν*_*re*_, and *ν*_*sn*_.

By taking the Fourier transform of Eqn. (8), we can represent the cortical excitatory field (*ϕ*_*e*_) in terms of the external sensory input field (*ϕ*_*n*_) in the Fourier domain:

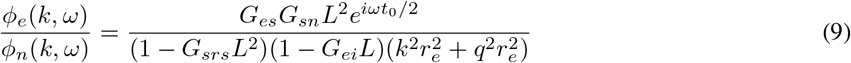

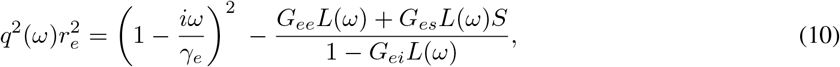

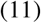

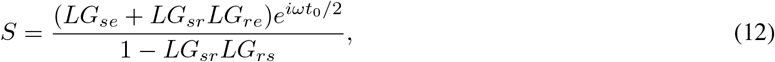

where *k* = 2*π/λ* is the wave vector with wavelength *λ*.

In a steady state, can assume *V*_*a*_ to be the only the perturbations to the function and take a linearized approximation of Eqn. (1), around the first term of the Taylor expansion. We define the parameter *ρ* as the derivative of the first term in this expansion. Hence we can reinterpret Eqn. (1) as:

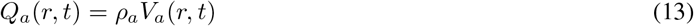

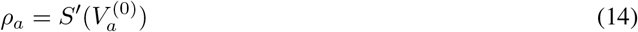

*G*_*ab*_ is defined as the gain value between populations *a* and *b*, describing the strength of the response in population *a* as the result of the unit input of population *b*, determined by all the scalars that affect the activation of a synaptic population:

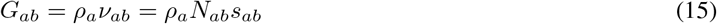

These gain variables are multiplicative, so that *G*_*abc*_ = *G*_*ab*_*G*_*bc*_. In this manner, the gains in functionally significant loops can be simplified as *G*_*ese*_ = *G*_*es*_*G*_*se*_ representing the excitatory cortico-thalamo-cortical loop directed by the thalamic relay nuclei, *G*_*esre*_ = *G*_*es*_*G*_*sr*_*G*_*re*_ representing the inhibitory cortico-thalamo-cortical loop directed by both the thalamic relay and reticular nuclei, and *G*_*srs*_ = *G*_*sr*_*G*_*rs*_ representing the gain in the inhibitory intrathalamic feedback loop.

In this case, we assume the uniform distribution of the cortical excitatory units (that is, spatially-uniform values of the wave vector *k*). If we approximate the brain as a finite-sized rectangular sheet with dimensions *lx* × *ly*, we can calculate the EEG power spectrum *P* (*ω*) as the integration of *ϕ*_*e*_(*k, ω*) over the wave vector *k*:

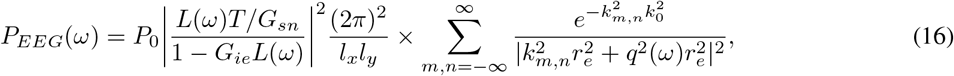

where:

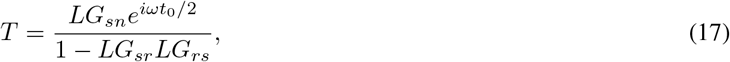

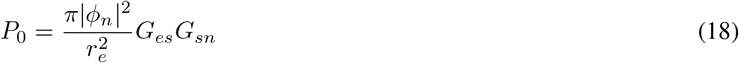

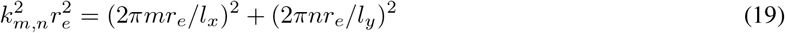

In Eqn. (16), the term 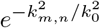 represents the low-pass spatial filtering induced by the dispersion of EEG electrical fields through the scalp and the cerebrospinal fluid between the cortex and the EEG sensor. This dispersion will also be spatially uniform given the uniformity of the vector *k*. The low-pass cutoff *k*_0_ is set at 10 m^−1^ based on prior empirical observations by Srinivasan et al. [85].

To mitigate the effects of high-frequency EMG artifacts introduced by pericranial, cervical, and extraocular muscles on the higher frequencies in of the power spectra [86, 87], an additional EMG power spectrum is added to that of the EEG [88, 89]:

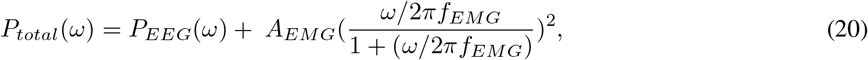

where the *A*_*EMG*_ term is fitted together with the other parameters for *f*_*EMG*_ in the range of 10 to 50 Hz.

It is worth noting regarding Eqn. (16) that by fitting this power spectrum function, we only get 5 gain (*G*) values (*G*_*ee*_, *G*_*ei*_, *G*_*ese*_, *G*_*esre*_, *G*_*srs*_) along with *α, β, t*_0_, and the fitted artifact term *A*_*EMG*_. There are infinite solutions to find *ν*_*a*_ with the power spectrum model fitting approach, since *ν*_*a*_ = *N*_*a*_*s*_*a*_. Similar to *ν*, the values of the gain parameters are negative for inhibitory synapses and positive for excitatory synapses, making *G*_*ei*_ and *G*_*sr*_ negative and all the other gains positive.

#### 2.3.1 *xyz* space

In the stable regions of the 9-dimensional parameter space at low frequencies, a reduced 3-dimensional space could be defined to represent the model parameters, in which: 1) *x* is the cortical loop gain and represents the corticocortical connection strength, 2) *y* is the corticothalamic gain and represents how effectively the thalamus can drive cortical activity, and 3) *z* is the intrathalamic gain. These three parameters are calculated via the following equations:

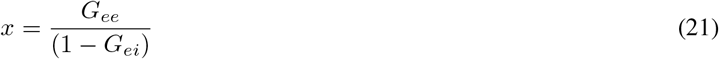

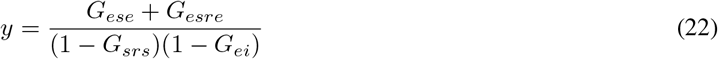

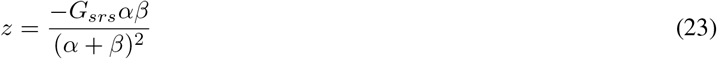

Each underlying state of brain activity gives more or less unique combinations of *xyz*. This system can be used to represent thalamocortical activity in many brain states with fewer dimensions than the entire fitted parameter set.

Eqn (22) asserts that the balance between cortical excitatory versus inhibitory activity determines the positive or negative sign of *y*. Excitation brings *y* toward more positive values, and inhibition will shift it to negative values. Eqns (21) and (23), respectively, indicate that the values of *x* and *z* will always be positive.

### 2.4 Simulation and model fitting

Simulations and model fitting were performed in MATLAB using the *Braintrak* library [89, 51, 90]. This toolbox implements a Markov Chain Monte Carlo (MCMC) method, using the Metropolis-Hastings algorithm for model fitting. The analytic power spectrum of the model, as defined in Eqns. (16) & 20, was fitted to the empirical power spectra from 30-second windows in the data. The parameters implemented were restricted to the stability limits defined in the previous literature [51] to ensure the biological feasibility of the attained fitted parameters. Furthermore, the value of the gain parameters for all connections was limited to 20 (|*G*_*ab*_| < 20) to reduce the sensitivity of the model to noise in the input.

#### 2.4.1 Fitting metrics

Using the method described above, we fitted the parameters of the corticothalamic model to empirical data. We used chi-squared (*χ*^2^) error for model optimization, calculated between the 1 and 45 Hz frequency bins. The optimisation target function aims to reduce the error *χ*^2^ between the empirical and model-generated power spectra. In this model fitting approach, the parameter space is firstly explored during a random walk with a length of 50,000 and with large step sizes, accepting the top points which get us to a region close to the target values. After this “burn-in” period, Braintrak takes smaller steps to approximate the ground truth more closely.

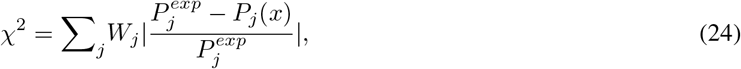

where *j* corresponds to each unique Fourier transform frequency bin of the power spectra. 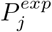 is the empirical (experimental) power spectrum and *P*_*j*_(*x*) is the predicted power spectrum for that bin. The term *W*_*j*_ is a scaling factor to increase the effect of the lower frequency bands compared to the higher frequency bands (*W*_*j*_ ∝ *f*^−^1), thus increasing the sensitivity of the optimizer to the high frequency bins of the power spectrum and minimizing its sensitivity to lower frequencies. This can be valuable in reducing artifacts observed in the EEG data, since the main artifacts affecting our 1-45 Hz window include the glossokinetic, movement, eye blink, and sweat artifacts, all of which produce low-frequency artifacts that must be mitigated [91, 92].

The complexity of the model was also calculated using the Akaike Information Criterion (AIC) [93]. AIC denotes the complexity of the combinations of parameters that together yield the model power spectrum. The lower the value of AIC, the simpler (or more parsimonious) the model. High values of AIC may denote overfitting of power spectra by fitting complicated combinations of parameters. AIC is described by this formula:

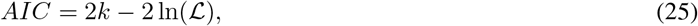

where *k* is the number of model parameters (9 in this case), and ℒ is the maximum of the likelihood function for this model.

### 2.5 Correlations between model parameters and health parame

By fitting the described data to this model across many subjects and over several datasets, we will be able to investigate the correlations between the changes in the model parameters and sleep stages, sleep quality metrics, and health markers that may have been collected from the subjects. For instance, the WSC dataset contains many such labels corresponding to many things such as the medications they were taking, their age, self-scoring surveys of depression and anxiety, and many markers of endocrine and metabolic health. We characterized characterized the differences between subjects on or off certain medications, and the changes in the model parameters between sleep stages. We further tested the existence of linear relationships between the changes in the model parameters and the health markers, using Pearson’s test via SciPy [94]. We then used the Benjamini-Hochberg method for False Detection Rate (FDR) correction [95].

Finally, we explored the ability of our estimated neurophysiological circuit parameters to predict disease outcomes using ML-based biomarker stratification. To this end, we separated the data into training and test groups, using a linear kernel Support Vector Machine (SVM) to classify binary health markers from the mean or variance of the fitted model parameters. The train/test data separation, load balancing, and training and testing of the model was done using the Python scientific computing library `scikit-learn` v1.1.2 [96].

## 3 Results

Our analyses in this study evaluated the methodology for neurophysiological modelling-based brain state estimation described above [51, 89] across several datasets recorded from research-grade and consumer-grade devices. In the following, we first summarize several key characteristics of the sleep EEG and hypnogram data used, and then turn to our model fitting results and their physiological interpretation.

### 3.1 Comparison of EEG features across sleep datasets

#### 3.1.1 Hypnogram-based sleep stage compositions

As can be seen in the group-averaged hypnogram summaries (Fig. 2E), sleep stages N1-N3 and W (wake) are well-sampled across all five of the studied datasets. REM sleep is also present in all datasets except Nap-EEG, since the 30-minute recordings used in that study are much shorter than the average 80-100 minute mark at which the first episode of REM appears [97]. The other four datasets all include several dozen minutes of REM-labelled sleep periods on average, although for the EDF-X dataset the average percentage of time spent in REM across the subject group is only 2.99% and the REM sleep latency (as seen in Fig. 2A), is unusually long (average of around 515 minutes).

**Figure 1:**
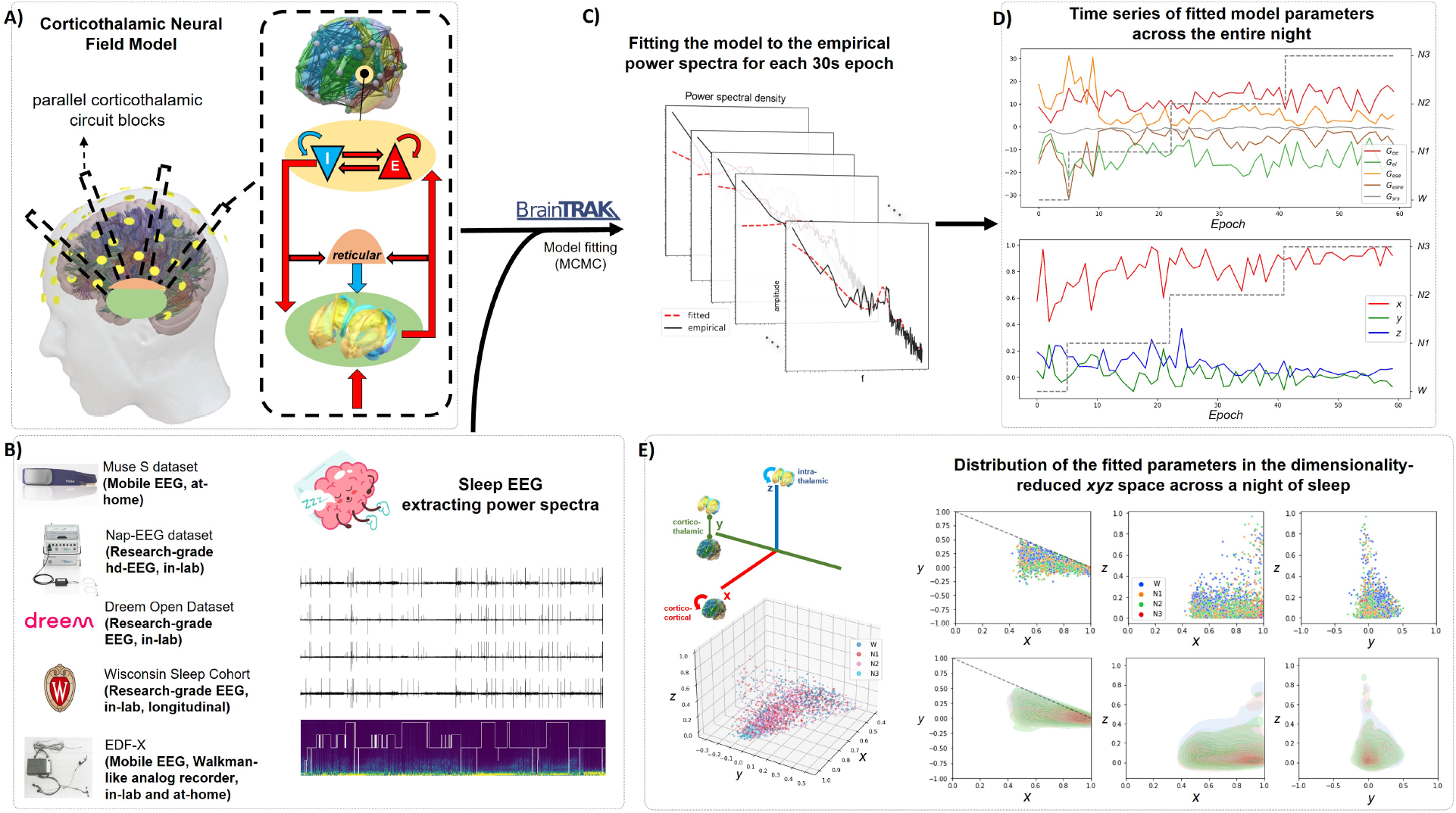
Method for fitting the sleep EEG power spectra to the model. **A)** Schematic of the neurophysiological model of the thalamocortical system, which simulates each channel of EEG as an independent active unit spanning the thalamic and cortical components. **B)** With this in mind, we accessed 5 datasets (EDF-X, Nap-EEG, DOD, WSC, MSD) which included sleep EEG and hypnograms. **C)** The *empirical* power spectra over 30-second epochs calculated using Welch’s method and the *fitted* power spectra generated using Braintrak to yield the fitted parameters. **D)** Time series of the fitted gain (*G*) parameters for subject 11, recording no. 1, from the Nap-EEG dataset. Each fitted epoch yields a set of 5 gains. The gain parameters are then used to calculate the circuit gain parameters *x, y, z*. **E)** Distribution of the parameters *x, y, z* across the entire Nap-EEG dataset is shown. Different sleep stages are denoted by the colour of the dots in the scatter plot. Different stages are clustered in different areas of the subspace. The dashed line in the 2D *x, y* plot marks the stability boundary at *x* + *y* = 1.

**Figure 2:**
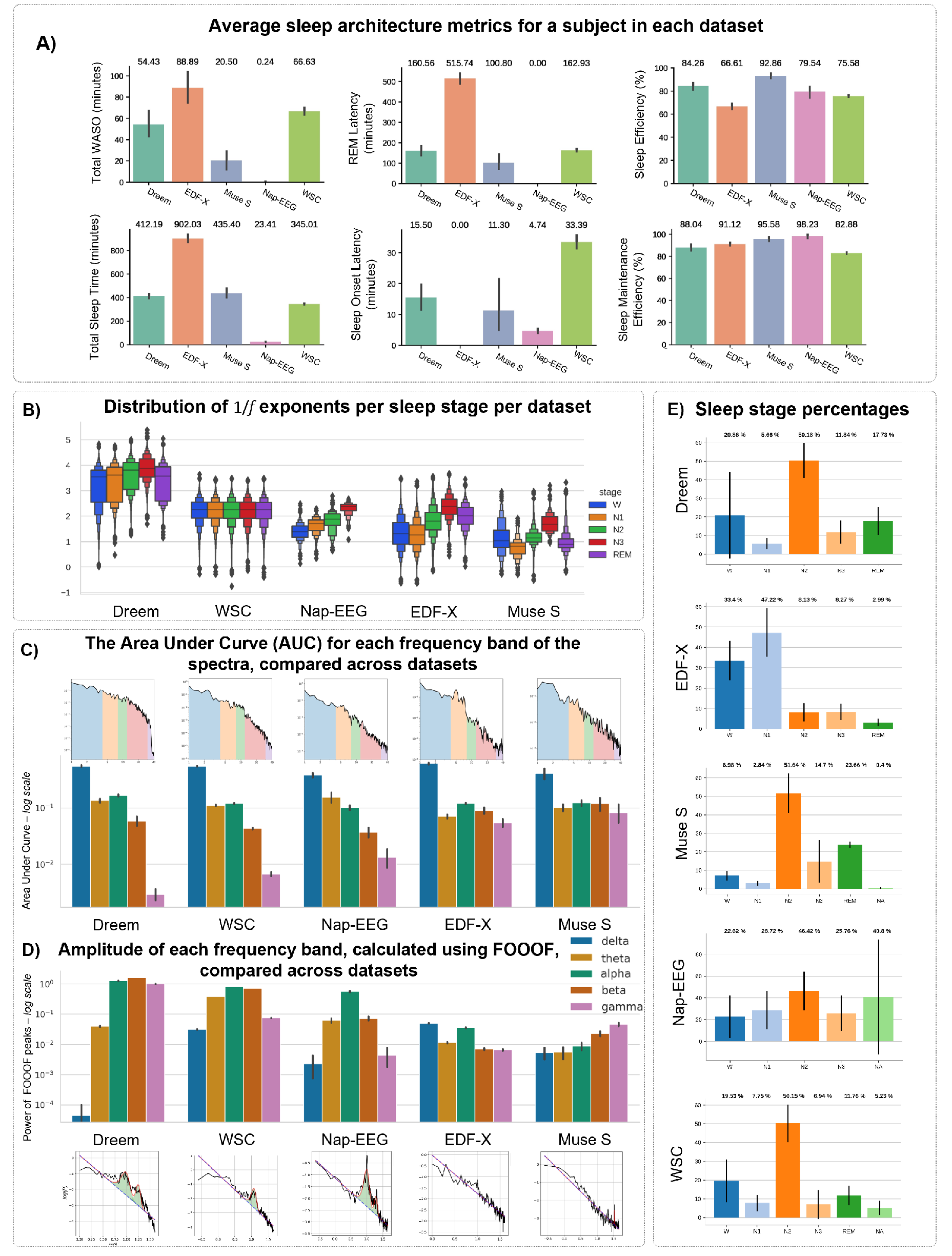
Comparison of sleep architecture metrics and power spectral features across datasets. **A)** Average metrics evaluating sleep architecture and sleep quality in a recording, compared across the datasets. Error bars represent the standard deviation of the values per subject. **B)** Distribution of 1*/f* exponents in each dataset, separated across the various sleep stages. **C)** *bottom:* Subject-averaged area under curve (AUC) of the EEG spectral power for each frequency band. *top:* Example power spectra from the dataset noted in the bar plot, with each under-curve band highlighted. **D)** Subject-averaged FOOOF-calculated peak power for each frequency band. **E)** Composition of the sleep stages forming the hypnograms in each dataset.

##### Improved sleep quality with mobile EEG

Sleep efficiency (SE, the percentage of time from the whole recording spent in sleep) between all datasets is comparable, averaging between 66.61% and 92.86% — although we note that for three of the five datasets, this value is below the recommended healthy range of 80-100% [98]. This value is the highest for the Muse S dataset with 92.86%. Sleep maintenance efficiency (SME, the percentage of time in sleep between the first and last stages of sleep) is also comparable for all datasets, with Nap-EEG performing the best among all (98.23%), and Muse S performing best for the whole-night recordings. For Muse S, this is most likely due to the improved comfort factor associated with the light and non-intrusive nature of mobile EEG headsets and the fact that the recordings are done at the subjects’ home and in their familiar and comfortable beds. This interpretation is further corroborated by comparing the subject-averaged total minutes of “Wake After Sleep Onset (WASO)” between the different datasets. Muse S subjects spend an average of 20.50 minutes awake after sleep onset, which is also the lowest among the whole-night recordings, further demonstrating that the subjects have less interrupted sleep when using mobile EEG equipment. This value is comparable to the range of 54-88 minutes for the three research-grade whole-night recordings (DOD, EDF-X, and WSC). Subjects also fall asleep faster with Muse S (average of 11.30 minutes Sleep Latency) than all other whole-night recordings, with the exception of EDF-X, for which an accurate sleep latency could not be calculated (see Supplementary Material section 3.1). Thus, mobile EEG can contain a more naturalistic and representative sample of physiological states and sleep stages in a full night of sleep than conventional research-grade EEG, and help us evaluate the normative trajectories of their changes in health and disease.

#### 3.1.2 EEG power spectral features across sleep stages

In the next stage, we used two different prevailing approaches for a quantitative comparison of the power spectra across datasets:

I. In one approach, the background scale-free 1*/f* activity is separated from the oscillatory activity with defined frequencies, and then the properties of each periodic & aperiodic component is examined. We used the Python library FOOOF for this task, as described in section 2.2.2.
II. The second approach is to compare the areas under curve (AUC) for the power spectral plots in each of those frequency bands. We used the trapezoidal integration method to calculate the AUC.

##### Aperiodic components vary across datasets and track sleep stages robustly

We separated the periodic and aperiodic components of the power spectra using FOOOF and calculated the exponent of the 1*/f* component for each power spectrum. The range of the 1*/f* exponents from different datasets are vastly different. Regardless of this variance, moving from lighter to deeper NREM sleep is generally associated with an increase in the value of the exponent and the value is then reduced again in the transition to REM sleep (Fig. 2A).

The scale-free changes in the slope of the 1*/f* component, which are thought to be results of background physiological processes and general brain activity patterns [99] can disproportionately increase the AUC in low-frequency domains, including delta (slow) and sub-delta (infra-slow) frequencies. We see this by comparing the power of each band as compared using FOOOF vs. AUC methods in Fig 2: The datasets Dreem and WSC, which possess the highest average 1*/f* exponents (as noted in Fig. 2A) demonstrate highest AUCs in the delta band (Fig. 2B), but by using FOOOF to separate the periodic & aperiodic components and examine the height of each unique peak apart from the contributions of 1*/f*, we observe that the delta band is the least dominant of all peaks 2C.

In fact, we see in that comparison that delta is the highest-powered band compared to all other bands in each dataset if we only use the AUCs, but separating the 1*/f* component using FOOOF relegates the rank of the delta peaks amplitudes to the last place in all cases. We further note that if we rely only on the AUCs, the alpha peaks are either the highest or the second-highest calculated peaks in all datasets with the exception of the Muse S. This demonstrates the importance of the contributions of the aperiodic component to the power of each frequency band in the power spectra and how it affects the traditional AUC methods for calculating band power. Solely observing the AUCs for each of those bands without this separation would have concluded a domination of low-frequency activity for all datasets, with minimal difference across the datasets, but FOOOF allows us to make that distinction between the footprints of each recording setup on band peak amplitudes.

In the MSD data, the peak height is lower than the other datasets for most bands (Fig. 2C). Separating the 1*/f* components from the raw power spectra in for this dataset almost completely reverses the order of the peaks with regards to frequency: the 1*/f* -separated peaks are highest for gamma and the lowest for delta, but the the AUCs are highest for delta and lowest for gamma. This points to the fact that for this dataset, the 1*/f* component is more dominant than the periodic component and that the 1*/f* exponent and is a robust feature separating the sleep stages and the physiological state of the subject, as evidenced by its strong variation across different sleep stages for Muse S (Fig. 2A), consistent with prior literature showing that most of the variation in individual sleep stages can be explained by 1*/f* components [36]. The 1*/f* exponents are one of the canonical features of our physiological model’s power spectra as well, as described in Robinson et al. [30], where the power law of the power spectra is a defining feature of the system, directly correlating with cortical gains and the primary oscillation frequencies, thus also making it a suitable criterion for tracking the activity of the corticothalamic circuitry.

### 3.2 Physiological modelling

Using the neurophysiological model of thalamic circuitry described in Abeysuriya et al. [51], Abeysuriya and Robinson [89], we fit the EEG power spectra across the various datasets. Despite the considerable difference in the properties of the EEG power spectra across these datasets, the model still performs well in fitting to all data. Goodness of fit was satisfactory, with all models demonstrating similar distributions of error (*χ*^2^) and model complexity (AIC), meaning that the model is fitting closely to power spectra without over-complicating the model parameters. The error distribution is marginally higher for Dreem than for other datasets which could be explained by the wide distribution of 1*/f* exponents in this dataset (Supplementary Fig. S1). The resulting fitted model parameters from all datasets exhibit patterns of change across sleep stages that are similar and in line with prior literature on thalamocortical communication in sleep. Corticothalamic communication is reduced in sleep and decreases further as the subject goes into deeper NREM sleep stages.

#### 3.2.1 Physiological transitions from light to deep NREM

The regular progression of sleep stages throughout a night of sleep commences with wakefulness (W), then transitioning to light NREM sleep (N1), followed by deeper stages of sleep (N2 and N3), eventually reaching REM sleep. Each individual cycles through the REM/NREM stages multiple times, with the cycles taking an average of 90 minutes [97]. In this transition within NREM sleep from N1 to N2 and to N3, the physiological properties of the functional brain circuits change along a clear trajectory [51], which we quantify using our neurophysiological modelling approach.

We first demonstrate this characteristic trajectory [51] and the differences between sleep stages using power spectra and fits from the Nap-EEG dataset. Similar overall results are obtained with the other four datasets, which are detailed further in the Supplementary Material Fig. S4. During the transition from W to N3, the 1*/f* exponent becomes larger and peaks in the alpha frequency band are reduced as deeper stages are reached (Fig. 3A). Two major patterns are observed in the neurophysiological model in conjunction with this change in the power spectra:

**Figure 3:**
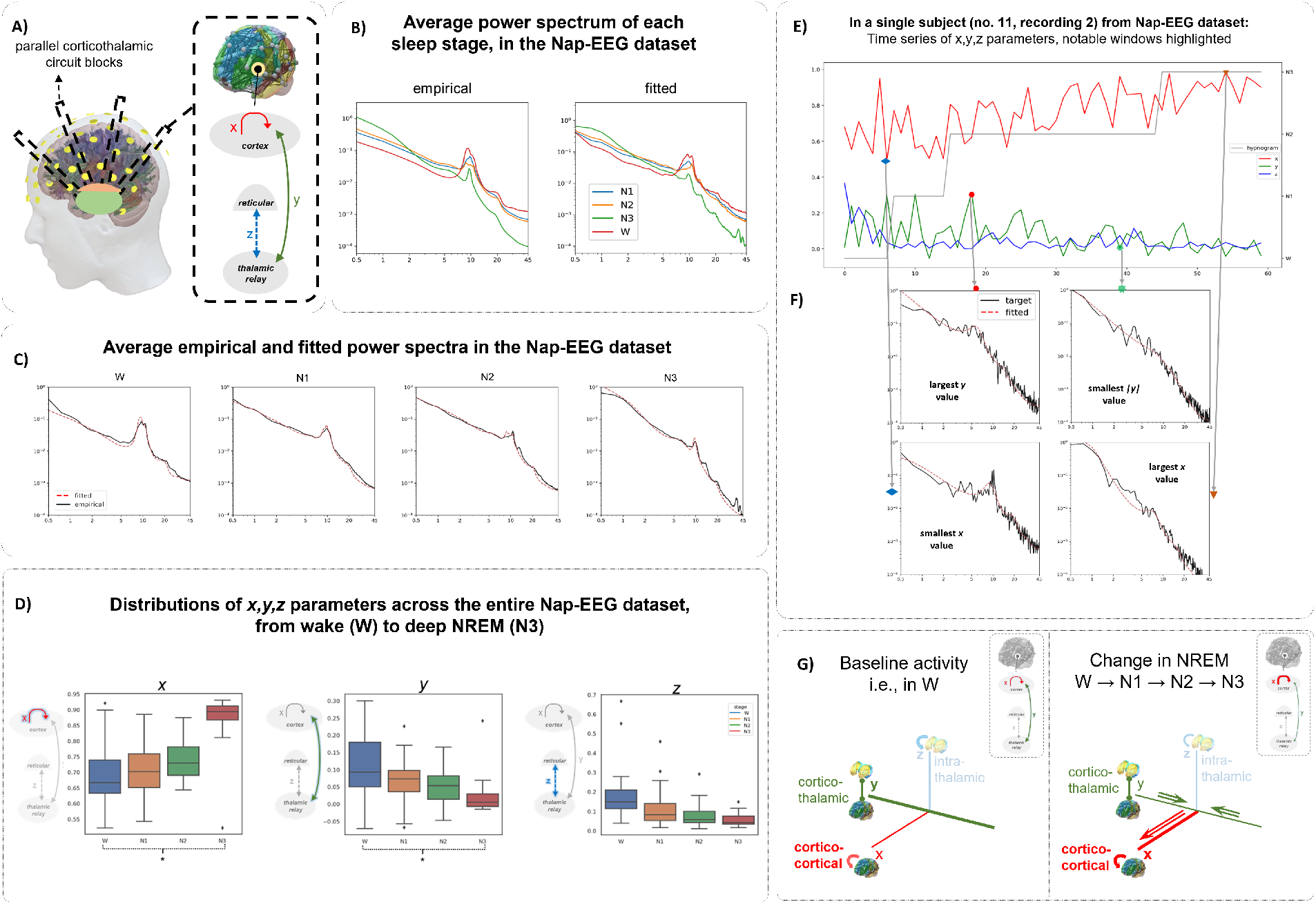
Tracking the changes in the parameters in different sleep stages in the Nap-EEG dataset. **A)** High-level schematic of the corticothalamic model, as described by the corticocortical (*x*), corticothalamic (*y*), and thalamothalamic (*z*) connection strengths. B) Average empirical vs. fitted spectra in the entire dataset, separated by sleep stages. **C)** Average power spectra in each sleep stage, separated between empirical and fitted. **D)** Box plots comparing the distribution of *x, y*, and *z* parameters across different sleep stages. Stages with significant difference in mean parameters are denoted by (*). **E)** Visualization of the *xyz* time series in conjunction with the hypnogram for one complete recording. **F)** Comparison of fitted and empirical power spectra at notable points in the whole-recording *xyz* time series with extreme *x* or *y* values, noting the associated alterations in the power spectra. **G)** Schematic demonstrating the change observed in the following panels. As the subject transitions from light to deep NREM, the connection strength in the corticocortical circuit is increased and the connection strength of the corticothalamic circuitry approaches zero.

##### Corticocortical amplification is increased from W to N3

Figure 3C shows the distribution of the corticocortical (*x*), corticothalamic (*y*), and intrathalamic (*z*) loop gain parameters across all epochs in the Nap-EEG dataset. The parameter *x*, calculated according to Eqn. 21, represents the net corticocortical excitatory connection strength. This parameter takes values between 0 and 1, with values close to 1 demonstrating highest degree of excitatory corticocortical amplification via the the excitatory projections connecting the various cortical regions and lowest corticocortical inhibition.

As can be seen, the progression from lighter to deeper sleep stages (N1 → N2 → N3) is associated with an increase in the average value of *x*, with estimates of this parameter in N3 approaching its maximum value of 1. This observation was confirmed statistically with an independent-samples *t*-test, which showed a significant increase in *x* from W to N3 (*t* = 17.29, *p* < 0.001). The corresponding comparison was also statistically significant in the other four datasets (Supplementary Fig. S4). This points to an association between deep NREM sleep and reduced thalamocortical drive of cortical dynamics. The reduction in the absolute value means that the bottom-up thalamocortical drive, in either the inhibitory or excitatory modes, is reduced in NREM sleep.

##### Bottom-up thalamocortical modulation is reduced from W to N3

During the transition from lighter to deeper stages of NREM, the distribution of *y* parameters–which indicates how strongly the thalamus drives cortical dynamics via thalamocortical projections–becomes narrower and more leptokurtic, with the absolute value of *y* decreasing and approaching 0. Based on the properties of the *y* circuit parameter, this signifies a reduction in the influence of thalamocortical gains (both inhibitory or excitatory). Per Eqn (22), positive values of *y* would denote the dominance of the excitatory part of the corticothalamic loop (*G*_*ese*_) over the inhibitory part (*G*_*esre*_), and hence net excitatory bottom-up stimulation thalamo-cortically. In contrast, negative values of *y* signify the dominance of the term *G*_*esre*_, where the inhibitory effect is due to GABAergic projections from the thalamic reticular nucleus. The negative value *y* therefore denotes a net inhibition applied to the cortex by the thalamus. As can be seen in the middle panel of Figure 3C the absolute value of *y* in N3 sleep approaches 0, signifying the absence of excitation or inhibition driven from the thalamus towards the cortex. Similarly to the previous section, we used the independent-samples *t*-test to compare the |*y*| values between stages W and N3, demonstrating a significant reduction in the parameter (*t* = −11.48, *p* < 0.001). Again, this effect was replicated across the other four datasets (Supplementary Fig. S4).

To further confirm that this relationship constitutes an ordinal trend across all four sleep stages, we assigned NREM sleep depth values of 0-3 to stages W-N3, and performed Pearson’s *r*-test with these and the lumped circuit gain parameters. This returned significant correlations between sleep depth and both the absolute corticothalamic circuit gain |*y*| (*r* = −0.226, *p* < 0.001) and corticocortical circuit gain *x* (*r* = 0.301, *p* = 9.83 × 10^−45^).

#### 3.2.2 Relationship of physiological circuit parameters to periodic and aperiodic EEG power spectrum features

We have demonstrated physiological model-based extraction of information on corticothalamic system state from windowed EEG power spectra across sleep stages and in multiple datasets. A key question that this analysis raises is “what features of the computed spectra contribute to the estimated physiological parameters”? As noted in Figure 2, different sleep stages have characteristic fingerprints across the periodic and aperiodic (1*/f*) components of the EEG power spectrum, that are generally consistent across all five datasets studied here. Given the evident associations of each sleep stages with the corticothalamic circuit activity parameters (Fig. 3) as well as the aperiodic components of the power spectra (Fig. 2A), we aimed to directly determine the interplay between the strength of various thalamocortical sub-circuits (gain (*G*) parameters) and the broadband power and 1*/f* exponents of the power spectra, along with a comparison of how each of these parameters relate to the changes in the power spectra.

To determine more precisely how the aperiodic components of the empirical power spectra give rise to the fitted physiological model parameters, we studied the correlation of these parameters across the entire Nap-EEG dataset with the slope and offset of the 1*/f* (Fig. 4A). This step was repeated for all datasets and all gain parameters in the Supplementary Figs. S5 – S9. We then examined further the contributions of isolated individual parameters to the power spectrum structure, by first initializing models at the estimated parameter values from a typical fitted epoch, and then systematically manipulating each gain (*G*) parameter, per Eqns. (16) –(19), observing changes in the model power spectra (Fig. 4B).

**Figure 4:**
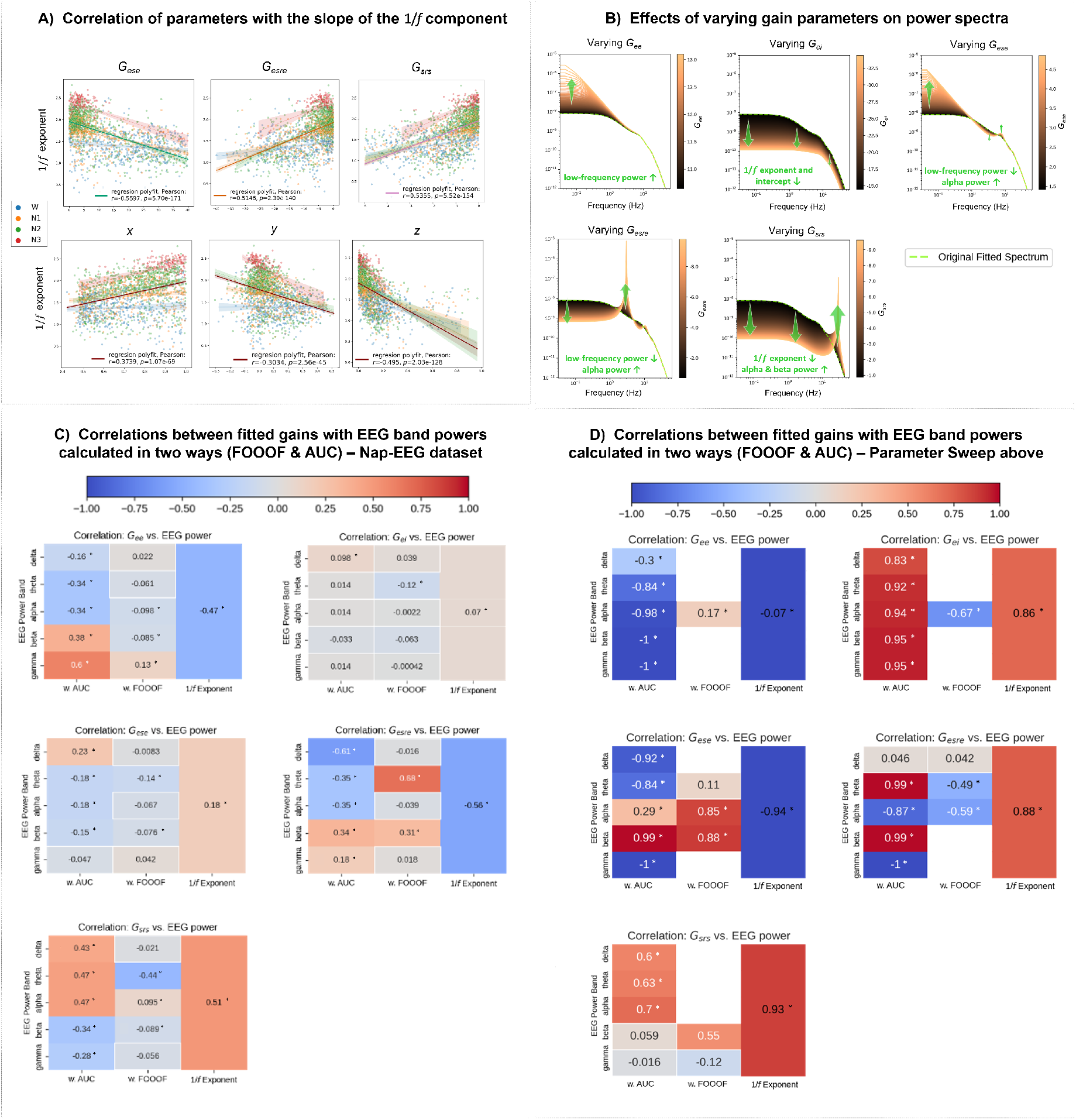
Correlation between model parameters and the shape of the power spectra. **A)** The correlation between the exponent of the aperiodic component with each thalamic gain parameter and *x, y, z*, calculated using Pearson’s *r*-test. *p*- and *r*-values reported in each legend. **B)** The effects of the incremental increase of each gain parameter on the shape of the power spectra. The baseline power spectra (in black) was taken from the fit to a real 30-second epoch (in Nap-EEG in stage N2 with no prominent peak). The absolute value of each gain value was increased in steps of 0.05 and the power spectrum generated from those parameters was generated (deducting 0.05 from negative gains and adding 0.05 to negative gains in each step. The colour bar denotes the values specified, starting from the darker color as the baseline and changing the gain parameters successively towards the lighter copper colour. **C)** Correlation of the gain parameters with the EEG Band power and the 1*/f* exponent in the Nap-EEG dataset **D)** Correlation of the gains with the power spectra resulting from the incremental sweep of the gain parameters.

##### Lower thalamic and corticothalamic gains generate a steeper 1/f

In all datasets, *G*_*ese*_—the circuit gain related to the positive feedback loop between the cortex and the thalamic relay nuclei—has a significant negative correlation with the exponent and the offset of the 1*/f* component (Figure 4A). That is, stronger excitatory corticothalamic feedback, signified by a higher *G*_*ese*_, results in a flatter 1*/f* component and a reduced area under the curve in the low-frequency domain. In general, we observe in the model fit results that epochs from sleep stage N3 tend to cluster in regions of parameter space with lower *G*_*ese*_ values and higher 1*/f* exponents. Thus, through the progression from wake to light and into deeper sleep, the exponent of the 1*/f* components increases, due to the progressive weakening of the excitatory corticothalamic feedback loop, as predicted analytically in Robinson et al. [30]. Concurrently with this, a positive correlation is observed between 1*/f* exponents and the negative-valued (inhibitory) gains of the loops associated with the thalamic reticular nucleus (*G*_*esre*_ and *G*_*srs*_). As can be seen clearly in Figure 4B), a decrease in the 1*/f* exponent and offset (flatter aperiodic component) is observed as both of these negative inhibitory gains become more pronounced. This is in line with observations by Abeysuriya et al. [51], in which a significant reduction in the strength of thalamothalamic inhibitory connections–represented by thalamo-thalamic circuit gain (*z*)–was observed in deeper sleep stages, which possess power spectra with higher 1*/f* exponents (Fig. 2A).

##### Greater corticocortical excitation can yield steeper 1/f components

The analyses in Figure 4B also show that amplifying the gains in the cortical excitatory (*G*_*ee*_) or inhibitory (*G*_*ei*_) connections results in a broadband increase in spectral power. The increase in *G*_*ee*_ is slightly more effective at increasing the lower frequency components. However, unlike the effects noted above for *G*_*ese*_, *G*_*esre*_ and *G*_*srs*_, modulation of *G*_*ee*_ or *G*_*ei*_ was not found to influence the observed spectra in this way in the datasets studied, and the correlations between these gain parameters and the 1*/f* components are not strongly correlated (Supplementary Fig. S7).

To further evaluate this effect of the gains on 1*/f* exponents, the analysis above was conducted for all the other datasets in addition to Nap-EEG (Supplementary Figs. S5–S9) which confirms this effect for most of the datasets. All of the datasets, with the exception of WSC, demonstrate significant moderate correlations between 1*/f* exponents and the corticothalamic gains (*G*_*ese*_ and *G*_*esre*_) and insignificant or significant mild correlations between cortical gains and the 1*/f* exponents. Abeysuriya et al. [51] report *G*_*ee*_ to highest variability amongst the gain parameters between different sleep stages. Our findings demonstrate that this is not mediated by the 1*/f* exponents at least in these datasets. In the case of WSC, the distribution of the 1*/f* exponents does not differ greatly between sleep stages as seen in Fig. 2A, which suggests that it might not be as adequate an indicator of the brain’s physiological state as it is for other datasets.

##### Fitted model parameters represent the power spectra robustly

In the next step, we examined the correlations between the fitted gain parameters and the power of each band and the 1*/f* exponent in the EEG power spectra, for each dataset and for the power spectra attained through the manual modification of single connection strengths. Pearson’s *r*-tests were implemented between the power of each band for each power spectrum and the gain parameters corresponding to it. The statistical significance of each correlation was corrected for the repeated hypothesis testing using the Bonferroni method [100] to prevent the false detection of patterns. It is observed that the offset & the exponents of the aperiodic components correlate with fitted parameters in the same direction and with similar strengths, as evidenced by Figs. S5 to S9. This analysis was completed for the Nap–EEG dataset in Fig. 3C and was repeated for all other datasets and all fitted and calculated parameters in the Supplementary Material Figs. S10 to S14.

It is notable that the gain parameters *G*_*ee*_ and *G*_*ese*_ are excitatory and hence have a positive sign. Meanwhile, the gains *G*_*ei*_, *G*_*esre*_, and *G*_*srs*_ are negative, and hence have a negative sign. Stronger connection strengths correspond to larger absolute values of these gains (more positive for the excitatory and more negative for the inhibitory). This must be taken into account when interpreting the correlations in Fig. 4.

Comparing the correlations for the fitted parameters (Fig. 4C) and the power spectra generated by changing the parameters (Fig. 4D) reveals that the positive gains are correlated and the negative gains are anti-correlated with the exponent of the 1*/f* component. The direction of the correlations is similar for both instances (the sign of the *r*-value). However, the intensity of the correlations (|*r*|) is larger for the manually-set parameters, which could be explained by the fact that all model parameters can also change for the fitted parameters, while in the manually-set instance, all parameters are fixed other than the changed parameter. The correlations between the changed parameter and the exponent in the set parameter instance is very strong (|*r*| *>* 0.86) for all parameters except for that of the cortical excitatory self-connection (*G*_*ee*_). Examining the parameters fitted to the Nap-EEG dataset, those correlations are moderate (0.5 < *r* < 0.6) for all parameters except for the cortical excitatory connection strength (*G*_*ee*_) and also the cortical inhibitory connection strength (*G*_*ei*_).

##### Physiological model captures changes in the spectra driven by both aperiodic and periodic components

As seen in Fig 4D, in examples such as the direct modification of the cortical inhibitory connection strength (*G*_*ei*_) alone, we observe a very strong correlation of the gain with the EEG band power calculated via the “area under curve” (AUC) method. But by separating the periodic and aperiodic components using FOOOF, we observe a different phenomenon; the aperiodic (1*/f*) exponent is increased in the same direction and approximately with the same intensity as the AUC band powers, but we only see one correlation with the FOOOF-discerned peaks in the alpha band, and no peaks were generated in any of the other EEG frequency bands. The case for the cortical excitatory connection strength (*G*_*ee*_) is somewhat different, where the AUCs of almost all bands except for delta have a very strong anti-correlation with the gain values (*r >* 0.84), but the FOOOF-discerned 1*/f* exponent does not show a notable correlation (*r* = 0.07) and the only correlating FOOOF-detected peak is alpha, with a weak correlation (*r* = 0.17).

As seen in Fig 4C, Comparing the FOOOF and AUC-detected phenomena in the parameters fitted to the Nap-EEG dataset demonstrates this common thread as well; all the connection strengths involving thalamus (*G*_*ese*_, *G*_*esre*_, and *G*_*srs*_) exhibit weak to moderate correlations with the AUCs of EEG bands, and they similarly correlate moderately with the 1*/f* exponents. Comparing the *r*-values of the correlations between FOOOF and AUC-measured peaks with these three gains demonstrates a strong disagreement between these two common metrics of power band estimation, where only two of the peaks are strongly detected by both methods, and in the case of theta band activity, they change in the opposite directions. This is due to the effects of the change in the 1*/f* component, where the exponent of this aperiodic component is correlated with the area under the low-frequency bands (e.g., delta and theta), and anti-correlated with the high-frequency bands. In other words, the shape of the power spectra and how it follows power law can change the detected values for those power bands.

Despite the variations, several consistent phenomena can be identified that are associated with higher absolute values of any of the three thalamic gains (|*G*_*{ese,esre,srs}*_|:

- The 1*/f* component is moderately decreased (−0.51 < *r* < −0.56).
- The AUC for the delta, theta, and alpha bands is decreased with a weak-to-moderate correlation (respectively,
- −0.43 < *r* < −0.61, −0.18 < *r* < −0.35, and −0.18 < *r* < −0.47).
- The AUC for beta and gamma band is weakly decreased (respectively, −0.34 < *r* < −0.38, −0.18 < *r* < −0.28).
- FOOOF peaks in theta are increased with a moderate correlation (−0.44 < *r* < −0.68).

We observe that the gain parameters attained by fitting the power spectrum can strongly capture the changes in both the periodic and aperiodic components of the power spectra. This model can describe both the broadband changes in the power spectra as described by the AUC measurements and the 1*/f* components, and the sharp peaks in narrower frequency bands, as measured via the FOOOF peaks. Among the fitted datasets, the 1*/f* exponents and the changes in the AUCs tend to describe the fitted gain parameters better than the sharp FOOOF-detected peaks in most of the datasets, as evidenced by Figs. 4C & 4D, pointing to the power of this model of EEG power spectra to represent the different power spectral features arising from the physiological properties of canonical brain circuitry.

### 3.3 Connecting the data-oriented and model-based observations for an interpretable understanding of sleep EEG

Features of sleep EEG change through health & disease, and by treatment. Common observations of such changes are usually data-oriented and describe statistical patterns in the time, frequency, phase, and spatial domains. Having described how the model tracks the periodic and aperiodic components of EEG power spectra, we now aim to see how these characteristics of the power spectra and their fitted model parameters map to various markers of sleep and mental health.

For that, we turn to two sources: **1)** The Muse S sleep EEG dataset includes repeated recordings from several users. We calculate key sleep quality scores from the hypnograms from these recordings to discern some nights with good or bad sleep quality. **2)** The Wisconsin Sleep Cohort dataset includes labels for many markers of physical and mental health, including medications taken by the participants.

We compare the power spectra and the fitted model parameters across different health states for the above two datasets to see find the changes associated with health, disease, and treatment.

#### 3.3.1 Observing changes in sleep EEG through repeated recordings using mobile EEG

To demonstrate the utility of mobile EEG for continuous monitoring of sleep EEG in repeated nights, we analyzed a set of repeated sleep EEG recordings from two users, who conducted sleep EEG recordings at least every other day in a span of 30-60 days. Using the hypnograms, we calculated key sleep quality scores via the Python library YASA [84]. Each user has their consistent range of values for these sleep quality metrics which can be persistent over multiple nights. Looking at the trends of these parameters over time, we can find patterns pointing to changes in the sleep quality.

##### Each user has a unique subspace of model parameters

repeated nightly recordings from the same user not only tell us about the night-by-night differences between their hypnograms, but also the large number of recordings allow us to characterize the subspace of parameters each individual will occupy in the available space of the model parameters. As we see in Fig. 5E, if we concatenate the parameters fitted to the epochs from all nights from each user, the general pattern of the inter-subject variability will start to be revealed. We can see that while the parameters of each sleep stage show great overlap especially around the mean values, the edges of these distributions are distinct, especially in wakefulness, N3, and REM. It is also worth noting that user 2 does not show N3 sleep in any of the nights. This shows the promise of mobile EEG for characterizing the normative sleep EEG of each individual based on their unique sleeping rhythm.

**Figure 5:**
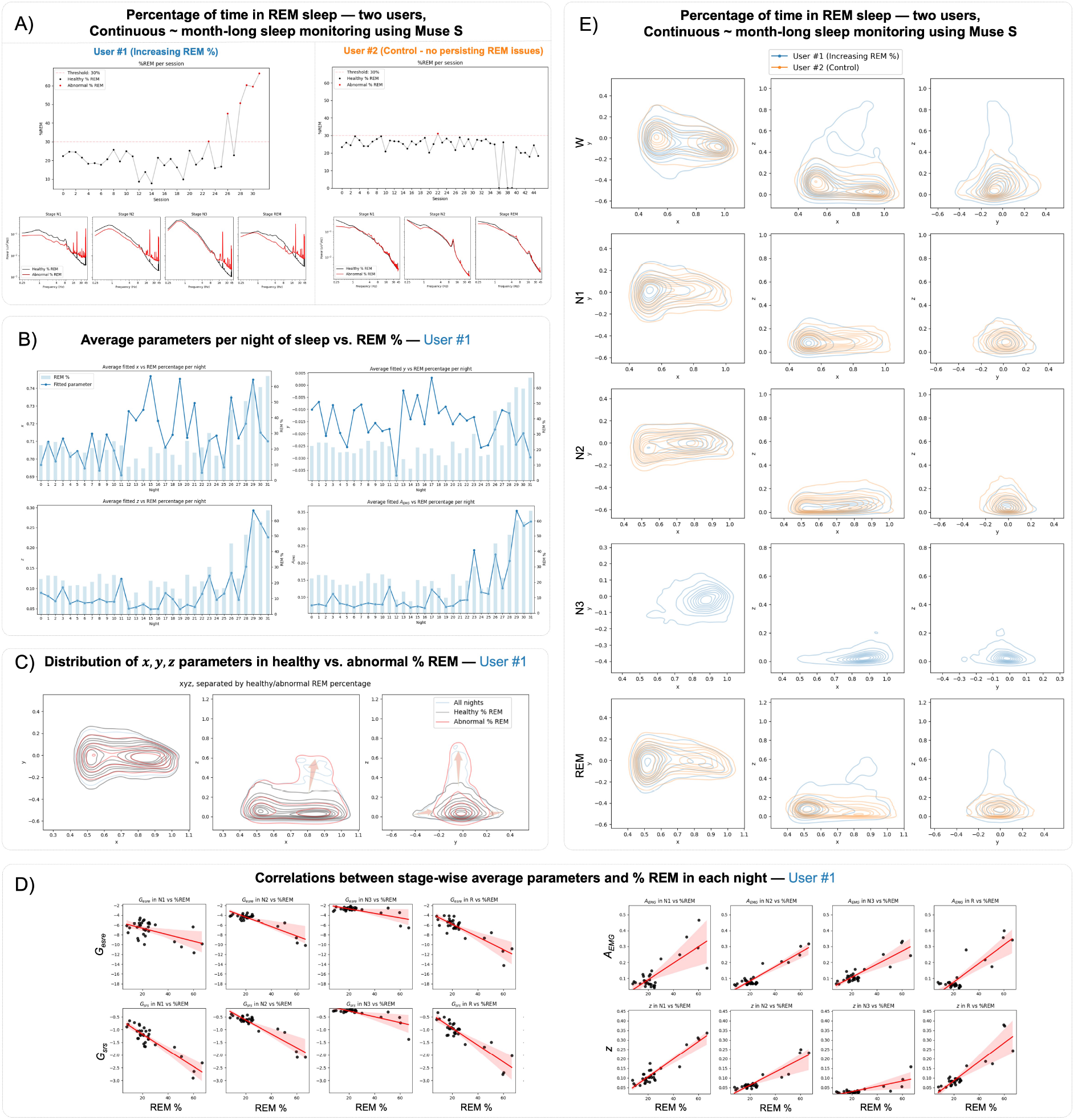
Sleep quality metrics in repeated nights. **A)** The percentage of time spent in REM sleep throughout the night for users 1 and 2 across the 30-60 day period. The stage-averaged power spectra are shown for nights with healthy vs. abnromal REM %. Values below 30% are considered normal in this case. **B)** Trends of the REM % (bars) vs. the trends of average *x, y, z* values (lines) for each recording, for user 1. **C)** The distribution of the *x, y, z* parameters across all epochs for healthy vs. abnormal REM % nights. Arrows point to the direction of the changes from healthy to unhealthy REM % nights. **D)** The average values of the thalamothalamic gain parameters for user 1, for nights with different REM % values. These trends are compared across sleep stages as well. **E)** The distribution of the *x, y, z* parameters across all nights for users 1 and 2.

##### One user shows a trend of increasing REM percentage and N1/N2 latency across consecutive nights

An interesting pattern that shows itself in the period of 30 recordings is that, as seen in Fig. 5, one participant (User 1) shows a trend of increasing REM percentage and increasing N1 & N2 latency over the observed period. Generally, such increase in the REM latency could be attributed to insomnia & sleep deprivation, or REM sleep disorders such as narcolepsy or REM sleep behavior disorder (RBD). But in the case of User 1, the Total Sleep Time (TST) is stable throughout the month, with the value of approximately 500 minutes, as are Sleep Efficiency (SE) & Sleep Maintenance Efficiency (SME) are also consistent, with respective ranges of 80-85 percent, and 80-90 percent. The REM latency for this user is fluctuating between 0 to 100 minutes, but this does not show a significant trend over time. This suggests that the increase in REM percentage is likely due to a disorder of REM sleep rather than a result of sleep deprivation or acute insomnia. In a systematic review, Boulos et al. [101] point to the parameter range of 8-21% for the average REM percentage across the age groups between 18 and 81 years old. With that in mind, we designated a threshold of 30% to denote if the REM percentage is abnormally high. User 1 shows a trend of increasing REM percentage with consistent repetitive at-home recordings using mobile EEG. User 2 does not show a significantly high number of sessions with high REM percentage, though this user does not show N3 sleep in any of the sessions. These frequent recordigns would have been difficult and expensive to maintain over such a long period of time. This demonstrates a practical utility of mobile EEG for long-term monitoring of sleep EEG to find these potential patterns of sleep quality deterioration. In this section, we compare the power spectra and fitted model parameters across these two users and across the nights with good vs. excessive REM percentage, to delineate a difference in the data and how underlying mechanisms can underlie these differences.

##### Nights of sleep with high REM percentage have lower 1/f slopes and higher thalamo-thalamic inhibition

In the next step, we compare the nights with high vs. low percentage of REM sleep. In figure 5A, we show that User 1 has a clear trend of increasing REM % up to values of around 60%. In the first 20 recordings, this user has a range of REM % between 10 to 25 %, and this range increases night after night to approximately 70% in the 31st recording. User 2 does not show a significant trend of increasing REM % over time. To understand if there effects of the potential case of disordered REM sleep, we separated all the power spectra from all epochs of sleep from nights with *healthy* vs. *abnormal* (excessive) REM percentage and calculated the average specta for each stage for both users.

We see in Fig. 5A that there are four clear changes in the power spectra from abnormal high-REM % nights compared to healthy REM % nights: 1) Descreasing 1*/f* slope (exponent) especially in N1 and REM. 2) Increased high-frequency power for all stages. 3) More prominent alpha peaks in N2, N3, and REM. 4) Increased high-frequency power (20 Hz and above) in all stages for high-REM nights. These differences do not seem to be salient for user 2 who does not show a significant trend of increasing REM % over time and only has one night just slightly above the REM percentage threshold of 30.

As we observe in Fig. 5B, the trends of model parameters across these recordings for user 2 show no significant trends in the corticocortical and thalamocortical circuit gains (*x* and *y*, respectively). But together with the increase in the REM percentage, the values of the thalamothalamic circuit gain (*z*) are increased. This increase suggests that higher thalamo-thalamic inhibition is associated with the boosted REM sleep throughout those nights.

To further examine if these average-level trends are mediated by the imbalance in the sleep stages, in Fig. 5D., we caculated the average parameter values per sleep stage per night and plotted them against the REM percentage for that night. The inhibitory circuit gains associated with the thalamic relay nucleus (*G*_*srs*_ and *G*_*ese*_) see a trend of increased inhibition (more negative values) in all sleep stages in the nights with higher REM percentage. This corresponds to an increase in the thalamo-thalamic circuit gain *z* in these nights. These correlations are most pronounced in N1 and REM sleep stages. We also see moderate correlations of increased excitation and inhibition across all other cortical and thalamic gain parameters (*G*_*ee*_, *G*_*ei*_, *G*_*ese*_), but those values cancel each other out in the overall circuit gains *x* and *y*, so they do not show a significant trend with REM percentage.

The distributions in Fig. 5C provide an overall view of these parameter distributions among the good vs. bad REM % nights. We see that nights with abnormal REM percentage have a different distributions in the *xyz* space, especially seen in the *xz* and *yz* orthographic projections, with extensions in the *z* direction towards larger values in unhealthy REM % nights.

##### Nights of sleep with high REM percentage have stronger high-frequency power

Another observable difference between the high-REM vs. low-REM nights is a clear increase in high-frequency power in the power spectra, as seen in Fig. 5A. As we argued earlier in this section, the combination of sleep quality parameters for this user points to a potential REM sleep disorder. In REM sleep behavior disorder (RBD) for instance, patients lose muscle atonia during REM sleep, which leads to them acting out their REM sleep mentations. Existence of an EMG rhythm in REM is the definitive diagnostic criteria for RBD [102, 103]. In this user, we have an increase in high-frequency power in high-REM nights, most prominently in REM sleep, which could be a result of muscle activity and EMG artifacts during sleep. In fact, in the mathematical model we use, there is an EMG term fitted to high-frequency power to mitigate the effects of EMG artifacts on the power spectra. The amplitude of this EMG term (the *A*_*EMG*_ parameter) is increased in the nights with higher REM percentage, as seen in Fig. 5B, further corroborating the suggested increase in muscle activity in high-REM nights.

#### 3.3.2 Model parameters are associated with markers of mental health

In this step, the fitted parameters of the Wisconsin Sleep Cohort (WSC) were analyzed in conjunction with the biomarkers included in this dataset. Using Pearson’s *r* test, the correlations between each of the 227 biomarkers and the average value per night for each of the 9 fitted model parameters were examined. To correct for the repeated pairwise correlation analysis, we used the False Discovery Rate (FDR) method introduced by Benjamini and Hochberg to correct the *p*-values in a ranked manner, taking into account the probable false positives in repeated testing [95, 104], bringing the *p*-value threshold for rejecting the null hypothesis from 0.05 to approximately 0.0396. There were 46 significant correlations between the parameters, but they were all weak–with the highest |*r*|-value for any correlation being 0.16. The significant correlations between these health labels and average model parameters per night can be found in 2. Despite the weak correlations, discernible patterns arise when observing which specific parameters correlate with which biomarkers. For example, the connection strengths in the inhibitory thalamothalamic feedback loop (*G*_*srs*_) and the the full thalamothalamic circuit (*z*) exhibit correlations with the administration of various medication groups and neurochemicals, such as alpha blockers, selective serotonin reuptake inhibitors (SSRIs), diabetes medication, and alcohol. This could be attributed to the various efferent cholinergic [105, 106] and serotoninergic [107] synapses that TRN receives, along with the complex calcium-dependent dynamics underlying its firing state and frequency [29], which would potentially be altered with the administration of these medications. The average thalamocortical circuit (*y*) exhibits significant correlations with biomarkers related to sleep quality and sleep debt, such as waking through sleep or daytime sleepiness. This is corroborated by evidence linking sleep deprivation to increased hyperexcitability and reduced specificity and functional connectivity in the thalamocortical connections [108, 109].

##### Selective Serotonin Reuptake Inhibitors (SSRIs) alter thalamocortical connectivity during deep NREM sleep

Given the importance of changes in sleep as a comorbidity of many mental health disorders, we next focused on studying the correlations of the average physiological circuit parameters fitted over one night with the WSC variables related to mental health–namely trait and state anxiety, scores from the Zung self-rating depression scale [110], and antidepressant medication. The Zung index is a normalized integer score value between 25 to 100, wherein the scores between 50 and 59 are scored as mild depression, between 60 and 69 as moderate depression, and any value higher than 70 as severe depression. Across our physiological model parameters we observed a weak but significant correlation (*r* = 0.10, *p* = 0.01) between the consumption of SSRI antidepressant medication and the gain of the thalamothalamic circuit (*z*).

We first tested whether the SSRI medication successfully reduces the severity of depression. In total, in 437 (78.60%) of the fitted recordings, the subject reported being on SSRI medication, and in the other 119 (21.40%), they were off SSRI medications. An independent-samples two-sided *t*-test demonstrates that the on-medication group has a significantly lower Zung index (*t* = −9.850, *p* < 0.001) than the off-medication group.

Next, we compared the composition of the parameters in different stages of sleep between the on- and off-medication group. We repeated the analysis in Fig. 3 on the on-SSRIs and off-SSRIs groups separately as well, to see if the transition from W to N3 yields the same reduction in |*y*| and increase in *x* in both subgroups. Independent *t*-tests were used to compare the means of *x* and |*y*| between the two stages W and N3. In the off-medication group, we see a significant *depth of sleep* effect–where the reduction in |*y*| is significant and negative (*t* = −7.778, *p* < 0.001) as is the increase in *x* (*t* = 2.004, *p* < 0.001). In the on-medication group, both of these effects are significantly reduced, with no significant reduction in N3 thalamocortical circuit gains (*t* = −7.778, *p* = 0.997) and very slight increase in the *x* values (*t* = 2.045, *p* = 0.020).

##### Interactions between depression or SSRI biomarkers with parameters x,y,z are nonlinear

We then attempted to see if the model parameters can be used to classify health labels directly, to test their potential standalone diagnostic use. We tried to predict whether the subject is on- or off-SSRI medication using the average or variance features of the WSC data, via classical machine learning approaches. We separated the data into training and test groups, with 80% of the fitted recordings in the training group and the other 20% in the testing group, with the on-SSRI group subsampled to match the size of the off-SSRI group. We then trained a linear kernel Support Vector Machine (SVM) to test if *xyz* in the on and off-medication groups are consistently separated using this support vector. The algorithm performed poorly at predicting the SSRI medication outcome. The linear-kernel SVM was not able to separate the two groups beyond chance level. This suggests that whole-night average parameter values are poor linear predictors of SSRI medication usage by themselves. For automated detection of the patterns observed in this paper using machine learning, we must utilize algorithms that can capture nonlinear relationships between the parameters and the health labels and the trajectories of change in the model parameters across a night of sleep, such as convolutional neural networks (CNNs), or apply dimensionality-reduction techniques such as singular value decomposition (SVD) or Principal Component Analysis (PCA).

## 4 Discussion

In this work we aimed to study how brain activity changes across different sleep stages, in health vs. disease, and as a function of recording technology (research-grade vs. mobile EEG). EEG power spectral density was calculated in 30s windows matching those of the hypnogram, delineating the frequency-domain characteristics of oscillatory and aperiodic background brain activity. Then, we used a neurophysiological modelling method introduced by Robinson et al. [45, 51, 89] to estimate various physiological parameters of corticothalamic brain circuits, and observe how these parameters change over sleep stages. Multiple sleep EEG datasets were employed to replicate our principal findings and to demonstrate the usage of this approach in various research and non-research scenarios, including most importantly, using at-home sleep EEG recordings from the consumer-grade sleep EEG headset Muse S. Changes in the 1*/f* -parameterization of the power spectra was shown to be significantly correlated with the corticothalamic gain parameters linked to bottom-up thalamocortical drive of the cortical activity, with the exponents becoming larger with depth of sleep (Fig. 4 and Figs. S5 to S9). Deeper NREM sleep stages were also observed to undergo a severance of effective bottom-up thalamocortical control, signified by reduced thalamocortical circuit gains (|*y*|) and increased cortical excitability, signified by elevated corticocortical circuit gain values (*x*) (Fig. 3 and Fig. S4). Administration of SSRI medication was observed to block this disintegration of corticothalamic connections in deep sleep. We additionally studied a case of an individual conducting repeated at-home sleep EEG recordings via Muse S, presenting with a REM parasomnia, associated with increased high-frequency EEG activity in the power spectrum, and an increase in thalamo-thalamic inhibition in the model parameter space. In summary, it was demonstrated that this physiological modelling approach can effectively integrate the periodic & aperiodic components of the EEG power spectra more robustly than common PSD analysis techniques and provide a reliable and physiologically explainable parameterization of those spectra in health & disease, and for the brain’s response to a treatment.

### 4.1 Key Results

#### Thalamic relay excitation increases the 1/f slope

A central result that was consistent across most of the analyzed datasets was that whereas connectivity strengths for cortico-cortical connections (*G*_*ee*_ and *G*_*ei*_, *x*) had negligible associations with the fitted 1*/f* offsets and exponents, strong correlations with 1*/f* features were seen for connections involving thalamic units (*G*_*ese*_, *G*_*esre*_, and *G*_*srs*_). The pattern is such that the higher the value of the gains (either excitatory or inhibitory), the bigger the exponent of the 1*/f* component, and so the more steep the background trend.

Previous literature points to the importance of the thalamic reticular nucleus as a regulator of excitatory thalamic nuclei activity, including relay nuclei [111, 29, 28]. This thalamic control loop attenuates the excitatory drive from thalamic relay nuclei to the cortex, thereby regulating the activity of the cortical neural populations underlying measured EEG signals.

As described in the 2 section, in the Robinson model, the thalamic relay nuclei (*s*) constitutes the main excitatory output of the thalamus, whose influence is balanced by the thalamic reticular nucleus (*r*) that implements a negative feedback loop, extinguishing the thalamic relay excitatory output. The loop gain *G*_*srs*_ gain parameter summarizes how the balance between these two units (*s* → *r* → *s*) enables this inhibitory feedback. One implication of our modelling results is that increased thalamo-thalamic inhibitory activity, signified by increases in inhibitory thalamic gains, flattens the 1*/f* spectrum by inhibiting the thalamocortical circuit driving the cortex. When this inhibition is removed, the network-level disinhibition in the cortex leads to a more steep 1*/f* slope. Previous authors (Gao et al. [20], Lombardi et al. [99] have suggested that higher 1*/f* exponents can be regarded as a criterion for higher cortical inhibition, driven not by thalamic but by cortical inhibitory populations, whereas our findings concentrate on the bottom-up thalamo-cortical axis of communication and how increases in its absolute gains lead to increased 1*/f* exponents.

A caveat for this model, as noted by Abeysuriya and Robinson [89], is that the gain (*G*) parameters are dependent on both the steady-state neural field and the synaptic strength of each population (per Eqn.(15)) and changes in either parameter can lead to a rise in the gain parameters, yielding infinite solutions for the exact delineation of these two parameters. Furthermore, another simplification in this model is the assumption of random outgoing synaptic connectivity, leading to *G*_*ei*_ = *G*_*ii*_ and *G*_*ie*_ = *G*_*ii*_, which may imbalance this cortical E/I balance estimation.

##### Thalamo-cortical disinhibition during the progression from wake to deep sleep

Using the neurophysiological model of thalamocortical system, we demonstrated that with the transition from wake to sleep, concurrent with an increase in the 1*/f* exponents, the values of the corticothalamic circuit (*x*) increase and the absolute values of the thalamo-cortical circuit gain (*y*) approach 0 (Fig. 3C). As we move from lighter sleep to deeper NREM sleep (from wakefulness to sleep stage N1, and then to N2, and then N3), the values of *x* increase further, such that in N3, their values are distributed very narrowly, close to the maximum value of 1.0. Concurrently, the value of |*y*| decreases, such that in N3, it has a narrow distribution close to 0 (both in Figure 4C). As noted, the values of *y* depend not only on the existence of thalamocortical activity, but also on the nature of its contribution (i.e. whether it influences inhibitory or excitatory activity from the thalamus to the cortex) [51]. In this sense, the deeper stages of NREM sleep, especially N3, involve increased cortical excitability, but at the same time, the thalamocortical population is insensitive to activity propagated through the thalamus. This highlights prior work by Nir et al. [40] demonstrating that EEG slow wave activity during deep NREM sleep is regionally and not globally synchronized, and that the oscillatory phases vary spatially over the cortical surface. Additionally, Massimini et al. [41] demonstrate that these slow waves can be locally interrupted and entrained using transcranial magnetic stimulation (TMS) at 1 Hz, suggesting a cortically-generated dynamic where local stimulation has the capacity to disrupt them. This pattern of dis-facilitation and dis-inhibition in N3 are in line with previous work using biophysical models denoting a reduction in thalamic excitation or inhibition in the brain in slow wave sleep [32] despite an increase in cortical synaptic strength [38, 112, 113]. Other work using neural mass models of thalamocortical circuitry by Müller et al. [114] has recently demonstrated the importance of the thalamus in maintaining the E/I balance in the cortex. They posit that adding a diffuse “one-to-all” connection term between the thalamus and the cortex, which is supported by empirical observations of the thalamic matrix nuclei helps recruit and dissolve the ensembles needed for cortical processing, and increases the transfer entropy from the thalamus to the cortex. They show that the effects of these matrix nuclei are highest in wakefulness and are decreased when modelling the effects of propofol anasthesia. This separation between conscious and unconscious states is in line with our observations regarding the bottom-up thalamocortical excitation or inhibition. Namely, we show that deeper sleep is correlated with the lack of large scale entrainment of the cortical activity by the thalamus, and the work by Müller et al. [114] delineates the other side of this same phenomenon that awake EEG corresponds to increases in the thalamocortical diffuse connectivity, driving the cortical activity from the bottom up. Further work utilizing the added thalamic nuclei in their work on the trajectories of activity in sleep can delineate the potential effects of matrix thalamus on sleep physiology as well.

##### The sign of y depends on the dataset, rather than the immediate power spectra

We found the mean and the mode of the fitted values for the circuit gains (*x, y*, and *z*) to be slightly different in various datasets. This could be justified by the different amount of time spent in different sleep stages (Fig. 2A) and thus the different oscillatory regimes dominating the data (Fig. 2C).

Despite the observation in Abeysuriya et al. [51] reporting excitatory thalamocortical regimes noted by positive *y* values in wakefulness decreases to negative values with the transition from wake to sleep, we observed that the numerical sign of *y* depends more on the datasets used than the stages of sleep. We noted the frequent “sign” of *y* to changes across datasets, irrelevant of wakefulness vs. sleep. For instance, 72.38% of all *y* values among 120,855 wake epochs in the EDF-X dataset were negative and 66.20% of all 2,119 epochs during sleep in the Nap-EEG dataset were positive. Hence, our work suggests a more nuanced take where the transition from wake to sleep shifts to a more inhibitory regime, marked by reductions in *y* along the depth of sleep axis, but that does not reflect an overall domination of bottom up inhibition as soon as sleep is initiated. Future work comparing the topography of these effects can shine light on the network-level variations in such changes.

##### Repeated at-home recordings using mobile EEG can help us better oberve parasomnias and modelling can help us understand the physiological basis of these conditions

We showed in this study an example of how repeated at-home recordings using mobile EEG can help us characterize the changes in EEG for a subject with a REM parasomnia. The repeated recordings allowed us to observe that the REM parasomnia is indeed consistent across nights and not a one-off sporadic event for the subject. For this subject, the EEG from the nights of high REM percentage (above 30%) had a flatter power spectrum, and the model parameters suggested a change in thalamothalamic gains in those nights.

The model seemed to fit more negative values for the gain parameters *G*_*esre*_ and *G*_*srs*_ in those nights. As observed in Fig. 4B, increasing these two gain parameters generate spectra with flatter 1*/f* components and higher alpha, beta, and high-frequency components. In other words, our model represents an increase in thalamo-thalamic inhibition with such a shape of the power spectra, and in the case of the parasomnias subject, has fitted power spectra with flatter 1*/f* components in the high-REM nights with higher thalamo-thalamic inhibition.

In this case, the increased high-frequency EEG activity in the REM parasomnia nights could be caused by a possible increase in the EMG-related artifacts in the EEG, as a result of increased muscle activity and loss of REM atonia, which is a characteristic of REM behaviour disorder. In this case, we also do not see all of the classic power spectral features of RBD, such as general slowing of the EEG, or general and widespread disruptions of N3 sleep. This is all further complicated by the great heterogeneity in the presentations of RBD for younger vs. older adults, and in the context of alpha-synucleinopathies [103, 115]. We highlight that these observations from one user are not complete, and nothing could be definitevely diagnosed without observing the subject’s EMG during REM sleep, which is the definitive diagnostic criterion. But this observed trend is promising for organizing focused future studies on REM parasomnias using mobile EEG from repeated recordings. In future work, the other actigraphy data that is already collected from many of the common consumer-grade sleep EEG headsets like Muse S can be combined with these power spectra to help with the diagnosis of REM parasomnias.

### 4.2 Limitations and Next steps

In this work we have focused primarily on changes in model parameters associated with transitions between sleep stages. However, these stages are far from the only physiologically-significant features we can extract from sleep EEG datasets. Other phenomena of interest could for instance be the dynamics of alpha activity in the final minutes of transitioning from wakefulness to sleep, which prior work has found to be associated with pathologies such as insomnia and sleep deprivation [116–118].

#### Studying sleep spindles

Another area of interest for future work that the framework presented here can be well suited to studying is transient oscillatory events in EEG traces such as sleep spindles and k-complexes. Abeysuriya et al. [49] utilize this model of the EEG power spectra to generate the power spectral density resulting from spindle generation. In this work the authors use stability analysis of the corticothalamic system to predict the nonlinear harmonic frequencies of the spindle peaks. In a follow-up study [50], they then demonstrated the existence of these spindle harmonics in empirical EEG data, as well robust fitting of an extended version of the model. In this case, the nonlinear harmonic frequencies of the spindle are resulting from the thalamo-thalamic feedback loop, and differ from the linear harmonic frequencies associated with the primary alpha and beta peaks. This study of the predicted harmonics in EEG power spectra can be useful for comparing the specific underlying corticothalamic connections generating such activity.

A long line of work on murine sleep EEG has demonstrated the importance of the corticothalamic system in generating sleep spindles. Empirically, shifts in the excitatory and inhibitory firing rates of various neural populations have been implicated in the generation of sleep spindles, initiated by a transition in the thalamic reticular nucleus [28, 119]. The circuit mechanisms and underlying mathematical structure of spindle generation in these detailed thalamic models [120] and the coarser-grained corticothalamic models [50, 49] may be related, but are not identical. They could be best understood as either complementary or competing candidate theories of this prominent phenomenon observed in human sleep EEG. An important direction for future work should be to compare and characterize the relevant parallels between these two frameworks, such as relating the models’ excitation and inhibition parameters to the 1*/f* EEG features across sleep stages described in this study.

The current work does not include any in-depth assessment of sleep spindles, as only one of the datasets used here (Nap-EEG) contained consistent spindle events with expert labelling in the EEG data. Separation of the spindles in the other datasets requires expert or machine learning-based detection of the spindles, which was out of the scope of the present work. Furthermore, sleep spindles start and end in short spans of approximately 2 seconds, which is much shorter than our standard power spectrum epochs of 30 seconds. The window sizes would therefore need to be substantially shortened to accommodate spindle-oriented analyses, which would in turn deleteriously increase the proportion of noise-driven peaks in the spectra, making model fitting less stable and consistent. In future work, we will study spindles as they appear in mobile EEG data specifically, and characterize changes in their occurrence, frequency, length, amplitude, etc ove sleep stages. These data can then be used to inform fitting of the corticothalamic model to the spindle PSDs, per [50, 49], thereby mapping these empirically-observed changes to transitions in the model parameters.

As we have indicated, sleep spindles are oscillatory events which begin, rise, decay, and then conclude in a well-characterized and parameterizable fashion. It is also notable that the phase and frequency of the spindles can vary spatially. In this study, due to computational limitations and variation in the data sets, we were restricted to fitting the power spectra to only one EEG channel. A logical next step would be to individually fit *all* EEG channels from the datasets, and analyze the parameters in the channel space, or to implement the spatial modes in the analytic power spectrum (*k*). The topography and spatial modes of these trajectories are topics of active interest in the field [121], and observing their changes in wakefulness and sleep in health and disease, especially in the context of mobile EEG, has clear scientific and clinical value.

In the future we will also consider transform-based machine learning, in which the transform is not merely a pre-processor but is also itself part of a neural network [122, 123], as well as phase-based methods [124]. Indeed, much of the important information in oscillatory activity during sleep is arguably better represented in terms of phase space, as well as scale space, phase scale, and the chirplet transform [122, 123], because sleep stages are often characterized by changes in frequency (acceleration of phase) [125–127]. Chirplet-based analytical approaches potentially offer a more biologically sympathetic perspective on neural signal analysis, which can aid corticothalamic modelling of sleep neurophysiology by better capturing time-varying frequency modulations in the EEG [128].

#### Implementation of this pipeline in large cohorts

Combining the above-outlined strategy with *de-novo* at-home sleep recordings using the Muse S headset, with a larger sample size than studied here, is a promising extension of the present work. In particular, this has major potential for studying sleep EEG features and the physiological underpinnings at-scale - both in terms of number of subjects (hundreds to thousands) and number of sleep sessions per subject (dozens or more). Adding biomarkers related to sleep quality and general health, for example through surveys or integration with other wearable biometric devices, would also be of great utility in delineating the physiological basis of those biomarkers.

In the present work, simple features of the distribution of 9 fitted corticothalamic model parameters across a night of sleep (such as mean, mode, and standard deviation) were used. In future work, using data-driven dimensionality reduction techniques to identify underlying sub-structures within these parameter values may prove an effective use of the physiological model outputs to help predict the health status and outcomes, both in extant datasets such as WSC [78], as well as new Muse S recordings with surveys described above.

### 4.3 Conclusions

In summary, our work has showcased the adaptability and reliability of this neurophysiological model [51, 89] for generating the trajectory of brain states during a sleep recording, utilizing a range of EEG data with various setups and recorded in various locales. This method adds another degree of physiological interpretability to the observations made based on EEG time series and power spectra. A robust interplay was observed between the aperiodic and periodic power spectral components, fitted model parameters, and their stage-dependent dynamics. This method can be effective for comparing sleep EEG between and among subjects and inferring latent health or disease states.

## 6 Acknowledgements

## Computing

The computing infrastructure for this work included the CAMH Specialized Computing Cluster (SCC) and the cluster Narval at Calcul Québec, part of the Digital Research Alliance of Canada. The SCC is funded by the Canada Foundation for Innovation’s (CFI) Research Hospital Fund. These platforms allowed for quick and robust large-scale model fitting to entire datasets and whole-dataset analyses.

## Author contributions

*(in alphabetic order)*: **CA**: Methodology, Resources, Data Curation, **DM**: Writing - Original Draft, Visualization, **JDG**: Conceptualization, Methodology, Software, Resources, Data Curation, Writing - Original Draft, Writing - Review & Editing, Visualization, Supervision, Project administration, Funding acquisition, **KK**: Writing - Review & Editing, **MAJ**: Methodology, Data Curation, **MPO**: Visualization, Writing - Review & Editing, **SB**: Writing - Review & Editing, **SH**: Writing - Original Draft, **SLH**: Supervision, Project administration, Funding acquisition, Writing - Review & Editing, **SM**: Funding acquisition, Writing - Review & Editing, **TM**: Conceptualization, Methodology, Software, Validation, Formal analysis, Investigation, Resources, Data Curation, Writing - Original Draft, Writing - Review & Editing, Visualization, **ZW**: Writing - Original Draft.

## Data and Resource Availability

1. The Sleep EDF–Extended (EDF-X) dataset version 1.0.0 used in this work is available on PhysioNet at https://physionet.org/content/sleep-edfx/, and described in Kemp et al. [70].
2. Dreem-Open-Datasets (DOD-H & DOD-O) and their annotations are available openly online. Full instructions on acquiring the data are included at https://github.com/Dreem-Organization/dreem-learning-open.
3. The Wisconsin Sleep Cohort (WSC) dataset is available on the National Sleep Research Resource (NSRR) [77] at https://sleepdata.org/datasets/wsc and can be accessed openly for academic research.
4. The Nap-EEG dataset is available via the Open Science Foundation (OSF) at https://osf.io/chav7/. Further information about the data is included by the authors at https://github.com/nmningmei/Get_Sleep_data.
5. The MCMC model fitting algorithm implemented on MATLAB is available at https://github.com/BrainDynamicsUSYD/braintrak.
6. The analysis and visualization code used in this paper is included in this GitHub Repository: https://github.com/GriffithsLab/MorshedzadehEtAl2024_sleep-eeg-nft.

