## Supplementary Material for "Corticothalamic modelling of sleep neurophysiology with applications to mobile EEG"

Morshedzadeh et al.

November 4, 2024

### S1 Overview of the supplementary material

In the supplementary material we provide further exploration of the model fitting quality with different parameters, and compare different model fitting factors and their effects on the final results. Next, we explore further the results observed in the main text. We demonstrate the analysis of all the dataset in this document—for instances where the main text figures only included one dataset for the sake of brevity.

In the supplementary methods we explore different MCMC chain lengths and the chi-squared error values (Figs. S1 and S2). In Fig. S4, the reduction in corticothalamic drive in deep NREM and the distributions of the reduced circuit gains $(x, y, z)$ is demonstrated for all 5 datasets. In Figs. S5 to S9, we expand on the effects of gain parameters on $1/f$ parameters.

### S2 Supplementary Methods

#### S2.1 Quality checks: Model Fitting

To further explore the performance of the fitting method, we made comparisons of the power spectra and parameters with different chi square values denoting good or poor fits, and also those generated utilizing different MCMC chain lengths.

##### *Effective fitting across most spectra, challenges with numerous peaks in a spectrum*

The fitted data uses the weighted chi-squared ($\chi^2$) difference between the generated and empirical power spectra to fit the model as described in Eqn. (24). The distribution of $\chi^2$ values features better fits with low error and poorer fits with higher error. In order to investigate what drives either case, we compared the power spectra separated by the percentile of $\chi^2$ errors for that epoch. Among the 149,403 epochs in the EDF-X dataset, the chi-square values ranged from 0.68 to

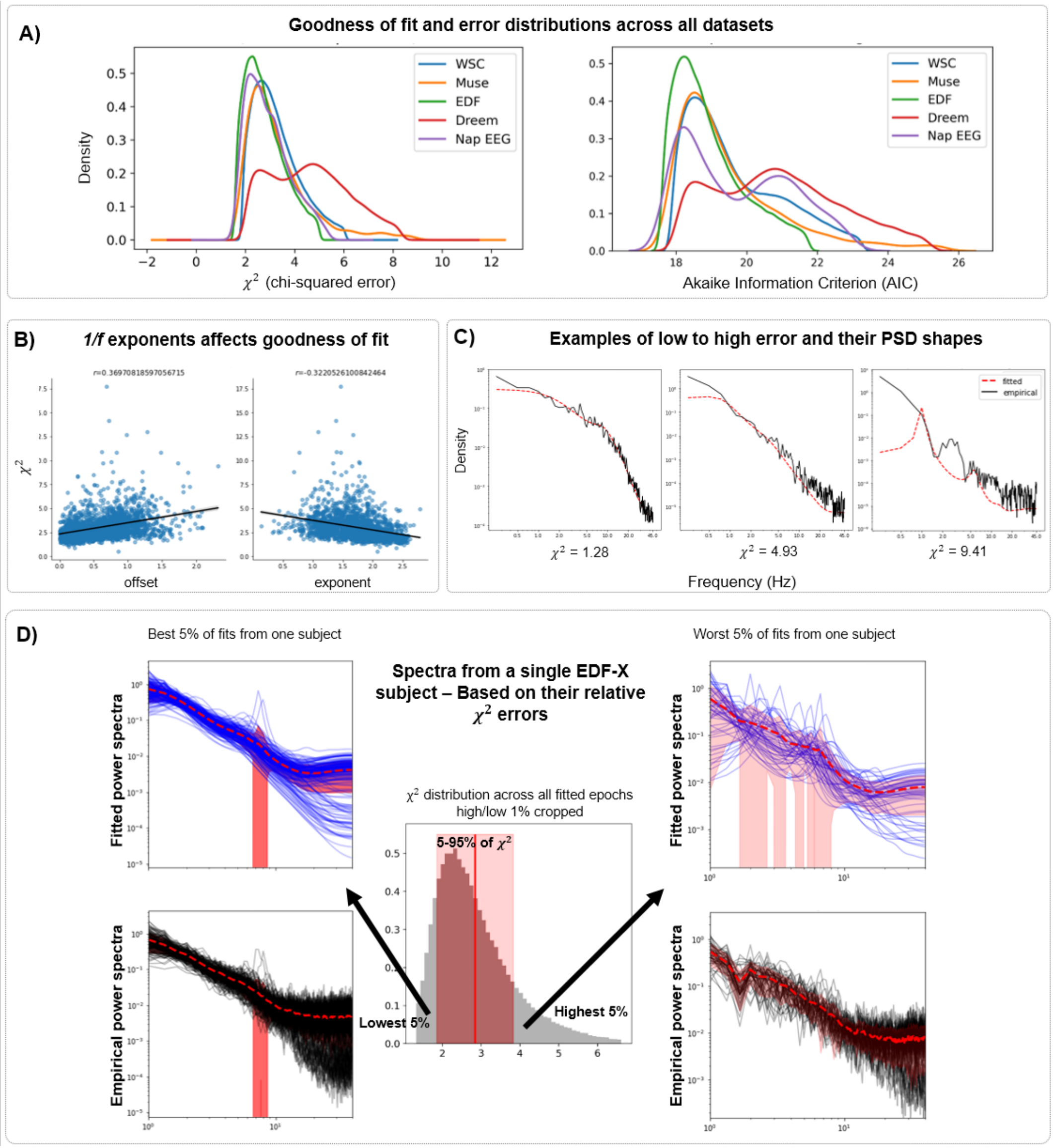


Figure S1: ***Comparing spectra with high vs. low $\chi^2$ errors in subject 1 of the EDF-X dataset.*** **A)** Distribution of the $\chi^2$ (chi-squared) errors and Akaike Information Criterion (AIC) from all spectra in each dataset. All datasets have similar distributions of errors and model complexity, which means that the model parameters are not over-fitting to the data. **B)** Generally, the model can fit better to power spectra with higher exponents and offsets closer to zero, noted by their correlations with the $\chi^2$ values **C)** Examples of empirical and fitted spectra with low, mid, and high $\chi^2$ errors. **D)** Errors from fitting to various power spectra taken from the distribution of $\chi^2$ errors from a single EDF-X dataset recording. Fitted (*top*) and empirical (*bottom*) power spectra demonstrated in each extreme of the high and low error distributions and the mean power spectra (dashed) are highlighted.

485, while 75% of all values were less than 3.37. The best 5% of fits in this dataset exhibit an lower average of 0.05 peaks per epoch and the poorest 5% of the peaks exhibit an higher average of 0.52 peaks per epoch, as calculated using FOOOF (described in section 2.2.2).

##### *Weak correlations between all model parameters and error values*

In the next step, the correlations of each of the fitted parameters with the chi-square values were calculated using Pearson's *r* test, in order to investigate whether the dynamics of any one specific parameter makes the model unstable and unable to fit the spectra.

Despite the significance of all correlation between the parameters and the chi-square error, all these correlations were weak (none yielded an *r*-value higher than 0.18, with 6 of 9 correlations with *r* values less than 0.10). In Figure S1, the comparison between power spectra in the highest 5% and the lowest 5% of $\chi^2$ values reveals that higher error values coincide with a larger high-frequency component. As described in (16), the neurophysiological model fits an EMG artifact term to the power spectra in order to improve the fits in this higher-frequency domain. The difference is not explained by the parameter $A_{EMG}$ either ($r = 0.07, p < 0.001$).

Our findings indicate that the Robinson power spectrum model excels in generating low-frequency and alpha-range peaks (and their harmonics), while also reproducing the $1/f$-like dynamics of the EEG robustly. However, the performance of this instantiation of the model tends to wane in the higher frequency domain since it lies beyond the primary time constants of the high-level thalamocortical circuitry within the model. This is particularly true in the absence of the spatial dispersion dynamics of the model implemented across different channels of EEG.

##### *Using different MCMC chain lengths does not meaningfully change the fits*

We explored whether using different chain lengths in the MCMC algorithm to generate samples from the probability distributions significantly changes the goodness of fits and the fitted parameters. Abeysuriya and Robinson [1] illustrate that using a chain length as low as 10,000 captures the dynamics of the model, as well as higher values such as 50000. To this end, we repeated the model fitting pipeline with chain lengths of 10,000, 20,000, 30,000, 40,000, 50,000 and 60,000, and compared the fitted parameters. While the time it takes to fit the power spectra increases with longer chain lengths, it does not change the chi-square values meaningfully. In addition, an approximately 25% change in the gain parameters is observed as a result of increasing the chain lengths and a change in the distributions of distribution of the synaptodendritic decay time ($\alpha$), synaptodendritic rise time ($\beta$), and thalamocortical characteristic propagation delay ($t_0$). These parameters are among those that determine the internal temporal dynamics of the model. As we increase the chain length, the bimodal distributions of the $\alpha$ and $\beta$ parameters seem to shift towards their lower peak, and the similarly bimodal distribution of $t_0$ shifts slightly to its higher values.

#### S2.2 Comparing model fits across datasets

The datasets used here were acquired from different sources. In total, 724,821 EEG windows of 30 seconds were fitted to the model. Out of these stages, the shares of each stage was as such: W: 159,808, N1: 113,732, N2: 298,871, N3: 51,075, REM: 74,874, unknown: 26,461.

##### S2.2.1 Fitted parameter distributions across datasets

We showed that the model can produce consistent fits across the five different sleep datasets used. We demonstrate here that mobile EEG, research-grade, and analog recording systems can produce fits that are bound and consistent

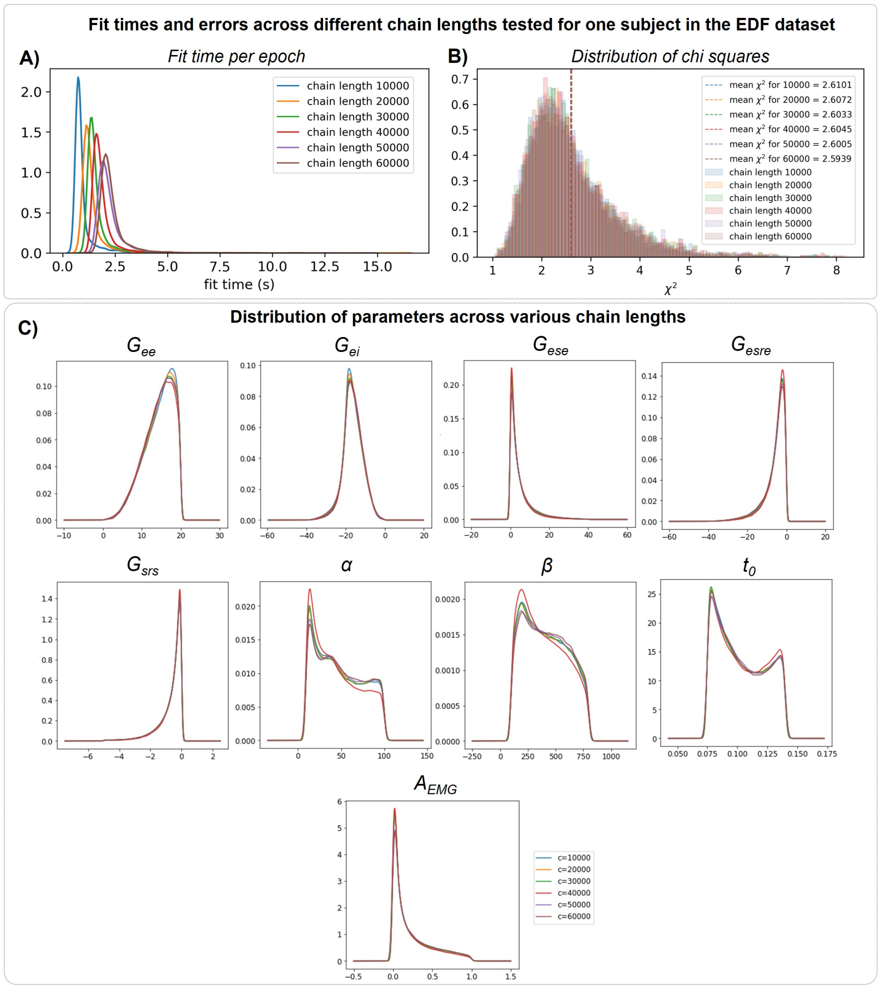


Figure S2: ***Comparing the parameters in the fits using multiple chain lengths/*** **A)** Distribution of model fitting times compared across different chain lengths. **B)** Distribution of $\chi^2$ errors utilizing the various chain lengths. **C)** Distributions of the fitted model parameters across different chain lengths.

using this platform. We demonstrate that even though different recording set-ups and pre-processing pipelines result in power spectra with different slope and peak properties, they can produce reliable results across hundreds of thousands of epochs.

Though, it is important to note that these differences in the power spectra lead to differences in the distributions of the model parameters across different datasets, and hence, the analyses on these model parameters must be done only between subjects with similar recording setups. For instance, Abeysuriya & Robinson [2] report that the values of $y$ tend to be positive among awake epochs and negative among epochs in sleep. In our work, we observe that the positivity or negativity of the $y$ parameter is more dependent on the dataset than on sleep stages. For instance, 72.38% of all $y$ values among 120,855 wake epochs in the EDF-X were negative, and 66.20% of 2,119 epochs in Nap-EEG in sleep were positive. We see this Figure S3K as well, where visibly, the colours of the dots (denoting sleep stage for each fitted epoch) in $y$ do not vary below or above zero.

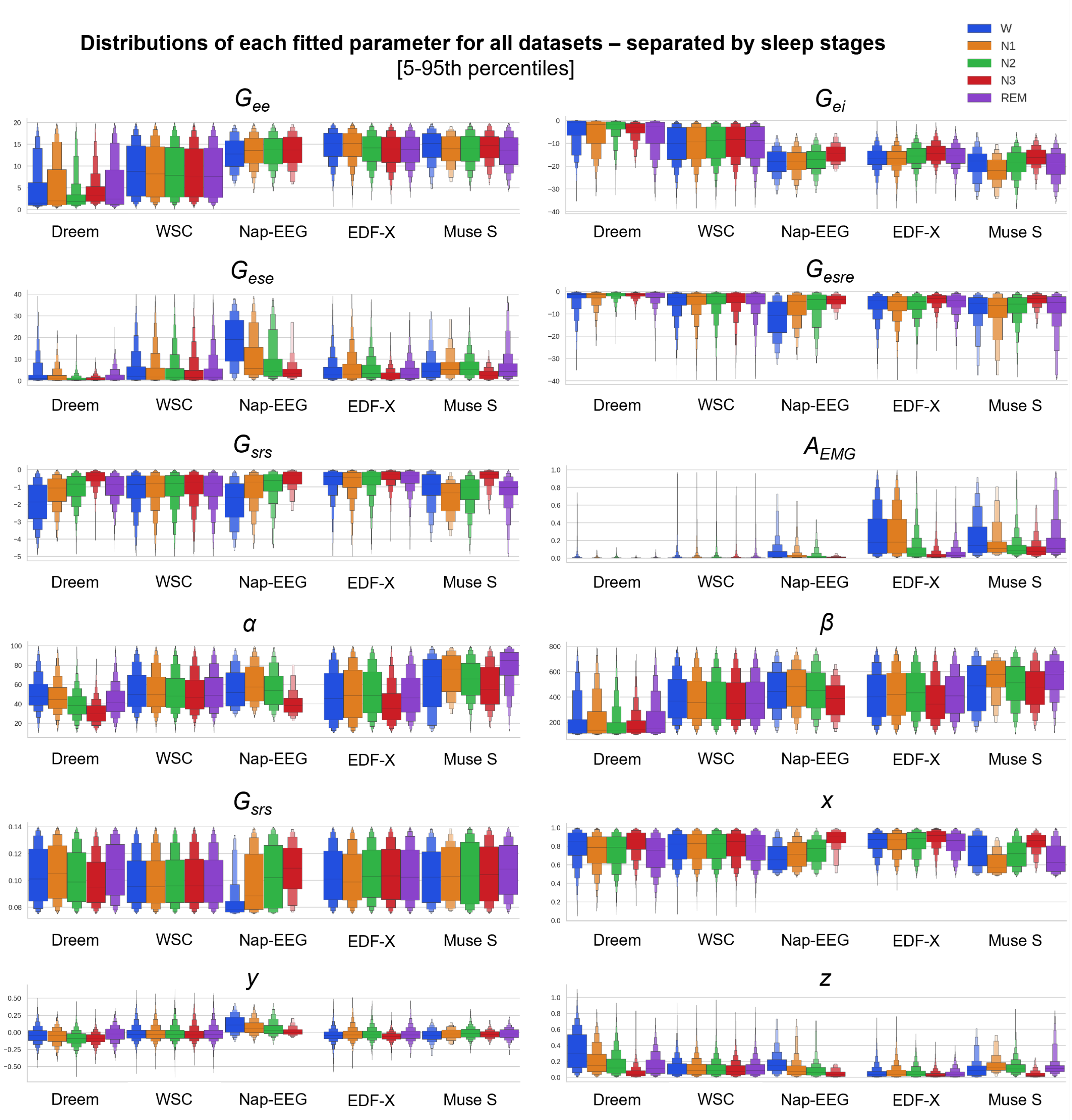


Figure S3: ***Boxen plots showing the comparison between fitted model parameter distributions across different datasets and sleep stages.*** In this figure, the parameter values for each of the 9 fitted parameters and the 3 calculated parameters $(x, y, z)$ are plotted. The 724,821 sets of fitted parameters separated based on the datasets from which they were acquired (sub-columns) and evident sleep stages during that epoch (differently coloured bars). Each rectangle of the boxenplot represents a *decile* (10×percentile) of the distribution. We observe differently clustering values for some of the parameters between datasets, but similar *differences* are seen between the sleep stages.

Table S1: Ranges of the model parameters in each dataset

| Parameter | Dreem | EDF-X | Muse S | Nap-EEG | WSC | **Abeysuriya (2016)** |
|---|---|---|---|---|---|---|
| $G_{ee}$ | 0.01 - 20 | 0.1 - 20 | 1.05 - 20 | 1.67 - 19.98 | 0.01 - 20 | 0 - 20 |
| $G_{ei}$ | -38.26 - 5.83 | -39.75 - 5.13 | -39.88 - 6.4 | -37.14 - 5.92 | -39.82 - 8.08 | -40 - 0 |
| $G_{ese}$ | 0 - 39.91 | 0 - 39.98 | 0 - 39.98 | 0.01 - 39.94 | 0 - 40 | 0 - 40 |
| $G_{esre}$ | -38.61 - 3.03 | -39.96 - 5.8 | -39.93 - 7.12 | -39.79 - 7.87 | -39.99 - 5.9 | -40 - 0 |
| $G_{srs}$ | -5 - 0.97 | -5 - 0.82 | -5 - 1 | -4.98 - 1.1 | -5 - 1.04 | -14 - 0 |
| $\alpha$ | 10.01 - 100 | 10 - 100 | 10.05 - 100 | 11.53 - 99.97 | 10 - 100 | 10 - 100 |
| $\beta$ | 100 - 799.79 | 100.01 - 799.99 | 101.69 - 799.71 | 100.98 - 798.41 | 100 - 800 | 100 - 800 |
| $A_{EMG}$ | 0 - 0.99 | 0 - 1 | 0 - 1 | 0 - 0.94 | 0 - 1 | 0 - 1 |
| $t_0$ | 0.02 - 0.14 | 0.02 - 0.14 | 0.02 - 0.14 | 0.02 - 0.14 | 0.02 - 0.14 | 75 - 140 |
| $x$ | 0.01 - 1 | 0.09 - 1 | 0.16 - 1 | 0.16 - 1 | 0.01 - 1 | 0 - 1 |
| $y$ | -0.8 - 0.67 | -0.66 - 0.58 | -0.65 - 0.46 | -0.3 - 0.51 | -0.86 - 0.81 | -0.8 - 0.8 |
| $z$ | 0 - 1.16 | 0 - 0.85 | 0 - 0.99 | 0 - 0.97 | 0 - 1.14 | 0 - 1 |

##### S2.3 Validation of the model parameters

In the next step, we compared the range of the parameters fitted to data from each set to the range of parameters from Abeysuriya and Robinson [1] to see if our fitted values are consistent and in line with the previously reported ranges in the literature using the same model. Table S1 highlights the ranges of the model parameters for each dataset to the values reported by Abeysuriya and Robinson [1]. The reported ranges are very similar across the various datasets and between our study and the previous literature.

#### S3 Supplementary Results

In this stage, we will review further additional detail regarding the analyses conducted on the various datasets in this paper, extending the results to all datasets.

##### S3.1 Hypnogram-based comparisons

In this supplementary section, we note some of the limitations and corrections implemented for specific datasets, based on their recording modes, which may affect some of the observations and calculations made for those datasets.

1. The Nap-EEG dataset includes only 30-minute naps, and all of the recordings are approximately 30 minutes in length. With such a setup, the subject cannot reach REM sleep, as noted in the main text.

2. The EDF-X dataset includes two subsets of recordings as described in Kemp et al. [3]: I) The *telemetry* subset that includes recordings around 9 hours in the clinic and II) the *cassette* that is around 20 hours long. There are many hours of wake data in the beginning that were cut down to near-zero, since it was hard to universally bring in a start-time that correlated with the typical *eyes-closed* state before sleep onset. This cropping step leads to an artificial "sleep onset latency" of near zero for this dataset, so we excluded EDF-X from the sleep onset latency Fig. 2. There still are long periods of wakefulness after the conclusion of the recording, which was kept, as it was relatively shorter in time. These added stages still lead to a higher representation of the stage "W" in EDF-X data compared to other datasets.

3. In all datasets, there are windows with "unknown" sleep stage labels. The data from those time windows are included in the hypnogram-based sleep architecture comparisons (section 3.1, Fig. 2A,E) and are excluded from further analyses since we were interested in changes in the parameters that are related to changes in sleep stages over a regular night of sleep.
4. The Muse S data is recorded from a light wearable device and the subject has increased potential of mobility and motility during sleep; in that light, we dropped the 30-second epochs with a voltage standard deviation higher that of the entire whole-night recording, leading to an approximately 7.48$\%$ decrease in the number of all epochs in this dataset. Similar to the EDF-X dataset, this correction was implemented after the hypnogram-based comparison step in Section 3.1 of the main text. So, the hypnogram-based comparisons in Fig. 2 include those dropped epochs as well. But the power spectral analysis (using AUC and FOOOF power band metrics) use the processed data post-dropping.

#### S3.2 Comparing N3 Corticothalamic effect across multiple datasets

In this step, we extend the analysis performed on the Nap-EEG dataset (Fig. 3) to all other datasets as well. The distributions of $x, y, z$ parameters are compared across various sleep stages, and their change from W to N1,2,3 is calculated via an independent samples $t$-test. The reduction in the absolute values of thalamocortical circuit gain ($y$) and increase in the corticocortical circuit gain ($x$) is similarly observed for all datasets (Fig. S4), as it does for Nap-EEG (3).

#### S3.3 Correlation of $1/f$ component parameters with the fitted gains

The correlation between the fitted gains in each dataset and the exponent and offset of the $1/f$ parameters were compared to investigate how these properties of the empirical PSDs affect model fitting. These gains signify the strength of connections among the different units of the model.

We show that there are significant and weak to moderate correlations between the exponents of $1/f$ and the cortical gains. But the correlations between the parameters and the thalamic gains tend to be more pronounced.

##### *Aperiodic components can drive the fits when the periodic components are reduced*

In each dataset, the correlation of aperiodic component parameters with the fitted parameters differs, based on the peakiness of the spectra in those datasets. In datasets that have smaller number of peaks per epoch (EDF-X with 0.15 peaks per epoch and Nap-EEG with 0.69 peaks per epoch), we see a significant correlation between the gains and the $1/f$ exponent and offsets. The correlations in these datasets tend to be stronger in the connections or circuits involving the thalamus (relay and reticular).Within these datasets with fewer pronounced peaks, the fitting is primarily driven by the $1/f$ component. In datasets with higher number of peaks per epoch (Muse, WSC, Dreem), weaker and less significant correlations between the aperiodic component parameters and the gains are observed.

##### *Steeper $1/f$ spectra generate reduced thalamic and thalamocortical gains*

In all datasets, $G_{ese}$ which is the gain related to the positive feedback loop between the cortex and the thalamic relay nuclei ($pyramidal \rightarrow thalamicrelay \rightarrow pyramidal$) has a significant and moderate negative correlation with the

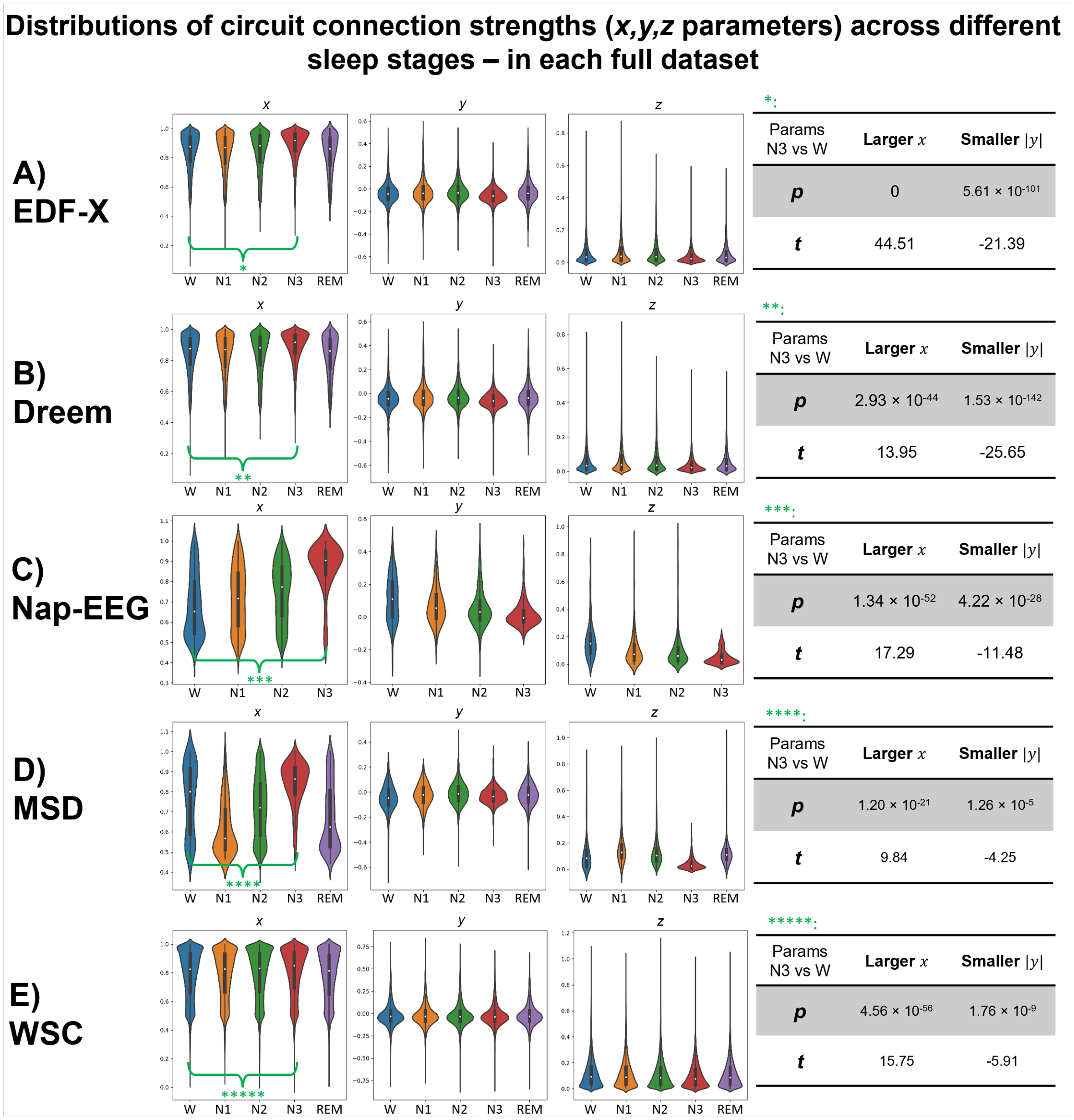


Figure S4: **Comparing the $x, y, z$ parameter distributions across different sleep stages, compared between all datasets**. In columns 1–3, the violin plots demonstrating the values of the $x$, $y$, and $z$ parameters across each sleep stage are plotted. In the 4th column, the $t$-tests comparing the mean connection strengths in the corticocortical ($x$) and the corticothalamic ($y$) circuits in wake (W) and non-REM stage 3 (N3) sleep are compared. Significant comparisons are denoted by green asterisks (*).

exponent and the offset of the $1/f$ component. In other words, stronger excitatory thalamocortical feedback loop, signified by a higher $G_{ese}$ results in a flatter $1/f$ component and a reduced area under the curve in the low-frequency domain. In almost all datasets, we observe the epochs from sleep stage N3 to cluster in regions with lower $G_{ese}$ values and high $1/f$ exponents. As we go into deeper sleep, the exponent of the $1/f$ components increase and with that, the excitatory thalamocortical feedback loop is weakened.

Simultaneously, a positive correlation is observed between the negative-valued (inhibitory) gains of the loops associated with the thalamic reticular nucleus ($G_{esre}$ and $G_{srs}$) and the $1/f$ exponents, pointing to a decrease in the $1/f$ exponent and offset (flatter aperiodic component) as the negative inhibitory gains become more pronounced.

For the spectra in the EDF-X dataset (Fig. S5) in which the average number of peaks per power spectrum is 0.15, a high correlation between the $1/f$ component parameters and the fitted model parameters is observed. More steep $1/f$ exponents (higher exponents and smaller offset values) increase the gains leading to cortical excitation ($G_{ee}$ and $G_{ese}$) and reduce the gains that lead to cortical inhibition $G_{ei}$, $G_{esre}$ and $G_{srs}$.

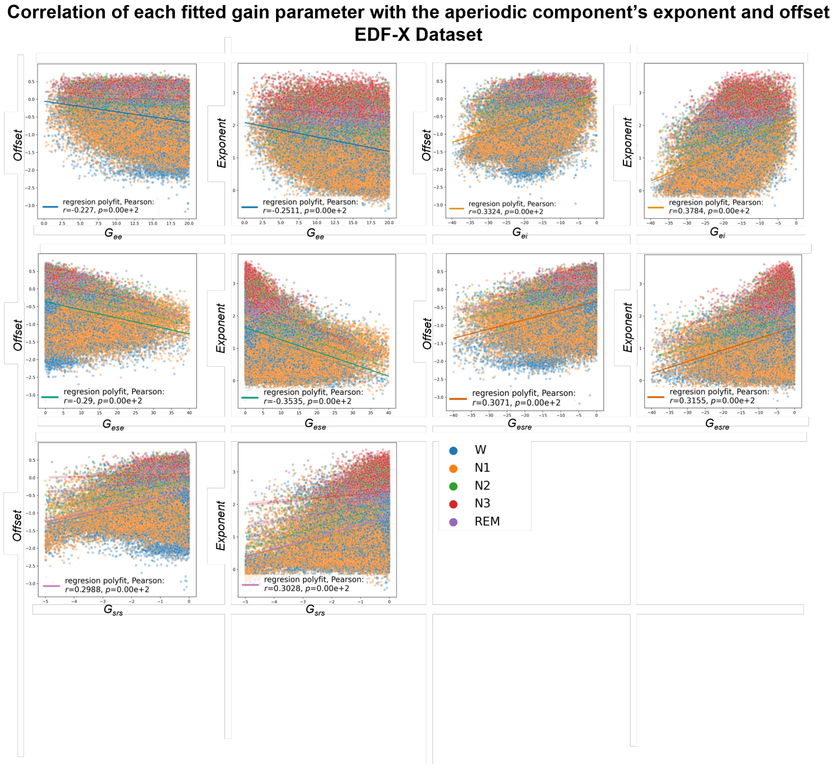


Figure S5: **Associations between gain parameters and $1/f$ components across the EDF-X dataset.** Points on each plot correspond to individual 30-second epochs throughout the entire dataset, color-coded by sleep stage: Wakefulness (W), NREM Stage 1 (N1), NREM Stage 2 (N2), NREM Stage 3 (N3), and REM sleep (REM). The odd columns from left present correlations with offset values, while the even columns depict correlations with exponent values. Each regression line is accompanied by a Pearson correlation coefficient ($r$) and a $p$-value, indicating the strength and statistical significance of the linear relationship.

In the Dreem dataset (Fig. S6), where the average number of peaks per power spectrum is 2.51, a weak-to-moderate correlation is observed between the $1/f$ components parameters and the fitted corticothalamic model parameters. The -positive- excitatory gains ($G_{ee}$ and $G_{ese}$) have an inverse correlation and the -negative- inhibitory gains ($G_{ei}$, $G_{esre}$ and $G_{srs}$) have an positive correlation with the $1/f$ exponent. This correlation is the weak for this dataset with the inhibitory thalamothalamic gain ($G_{srs}$) and moderate for the others. In general, the higher the absolute value of the gains (inhibitory or excitatory), the flatter the $1/f$ component.

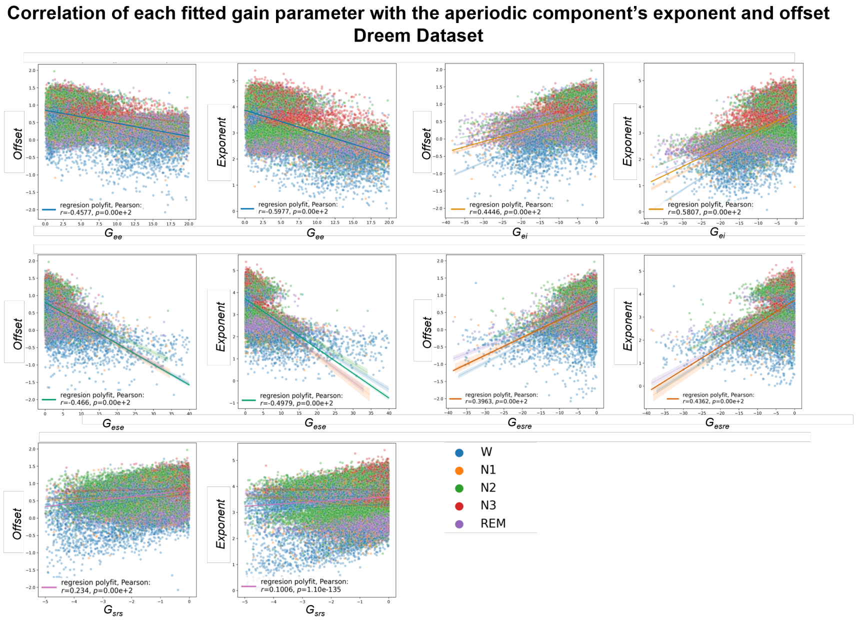


Figure S6: **Associations between gain parameters and $1/f$ components across the Dreem dataset.** Points on each plot correspond to individual 30-second epochs throughout the entire dataset, color-coded by sleep stage: Wakefulness (W), NREM Stage 1 (N1), NREM Stage 2 (N2), NREM Stage 3 (N3), and REM sleep (REM). The odd columns from left present correlations with offset values, while the even columns depict correlations with exponent values. Each regression line is accompanied by a Pearson correlation coefficient ($r$) and a $p$-value, indicating the strength and statistical significance of the linear relationship.

In the Nap-EEG dataset, the cortical gains ($G_{ee}$ and $G_{ei}$) seem to be less correlated with the $1/f$ component parameters, but the thalamic gains are more significantly and more strongly correlated with the gain parameters. Among the gains involving the thalamic relay or reticular nuclei, the the positive excitatory gain ($G_{ese}$) has an inverse correlation and the negative inhibitory gains ($G_{esre}$ and $G_{srs}$) have a positive correlation with the $1/f$ component. Among the fits from this dataset, the higher the magnitude of the gains, the flatter the power spectrum $1/f$ component.

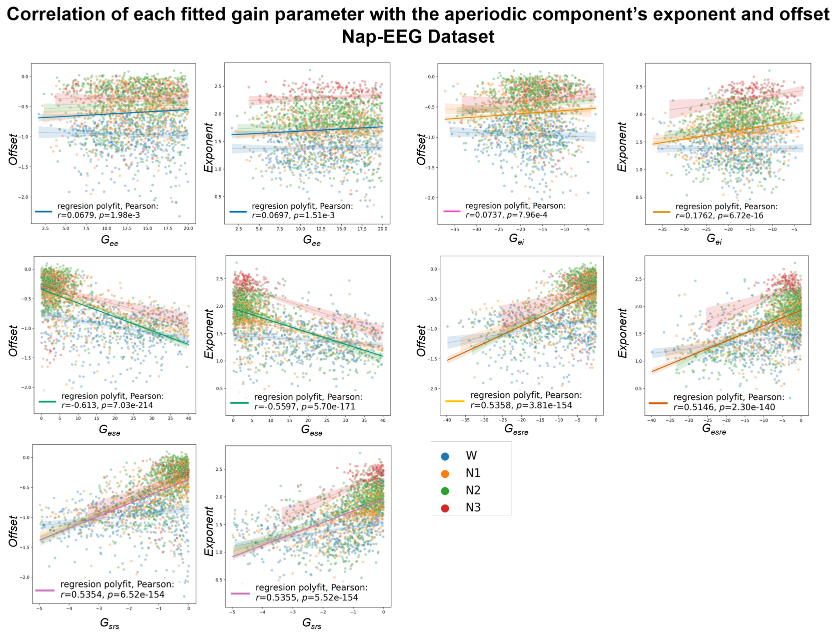


Figure S7: **Associations between gain parameters and $1/f$ components across the Nap-EEG dataset.** Points on each plot correspond to individual 30-second epochs throughout the entire dataset, color-coded by sleep stage: Wakefulness (W), NREM Stage 1 (N1), NREM Stage 2 (N2), NREM Stage 3 (N3), and REM sleep (REM). The odd columns from left present correlations with offset values, while the even columns depict correlations with exponent values. Each regression line is accompanied by a Pearson correlation coefficient ($r$) and a $p$-value, indicating the strength and statistical significance of the linear relationship.

For the Muse S dataset, the $1/f$ parameters are not significantly separated across the different sleep stages, noted by the non-separation of the differently-coloured markers. Most $1/f$ exponents and offsets cluster around similar values, with the value of 2 for exponents and -10 for the offsets. There is a significant and weak-to-moderate correlation, noted by $r$ values of 0.30 or less between the gain parameters and the $1/f$ component properties.

**Correlation of each fitted gain parameter with the aperiodic component's exponent and offset**
**Muse S Dataset**

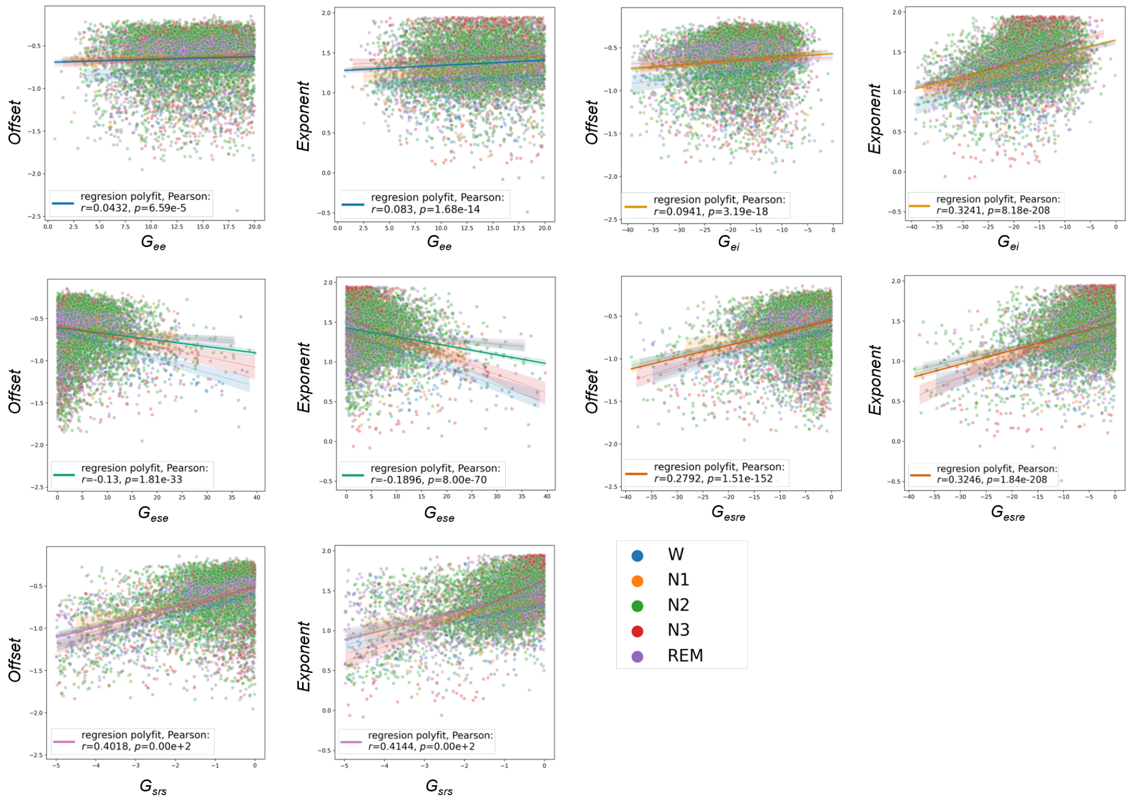


Figure S8: **Associations between gain parameters and $1/f$ components across the MSD.** Points on each plot correspond to individual 30-second epochs throughout the entire dataset, color-coded by sleep stage: Wakefulness (W), NREM Stage 1 (N1), NREM Stage 2 (N2), NREM Stage 3 (N3), and REM sleep (REM). The odd columns from left present correlations with offset values, while the even columns depict correlations with exponent values. Each regression line is accompanied by a Pearson correlation coefficient ($r$) and a $p$-value, indicating the strength and statistical significance of the linear relationship.

The distributions of $1/f$ component parameters in the WSC dataset have a very sparse distribution and are not separated as strongly across various sleep stages. Hence, the scale-free $1/f$ component in WSC is not as strongly the driving factor for the fits as the other datasets. The $p$ values of all correlations are high and the $r$ values are all smaller than 0.16.

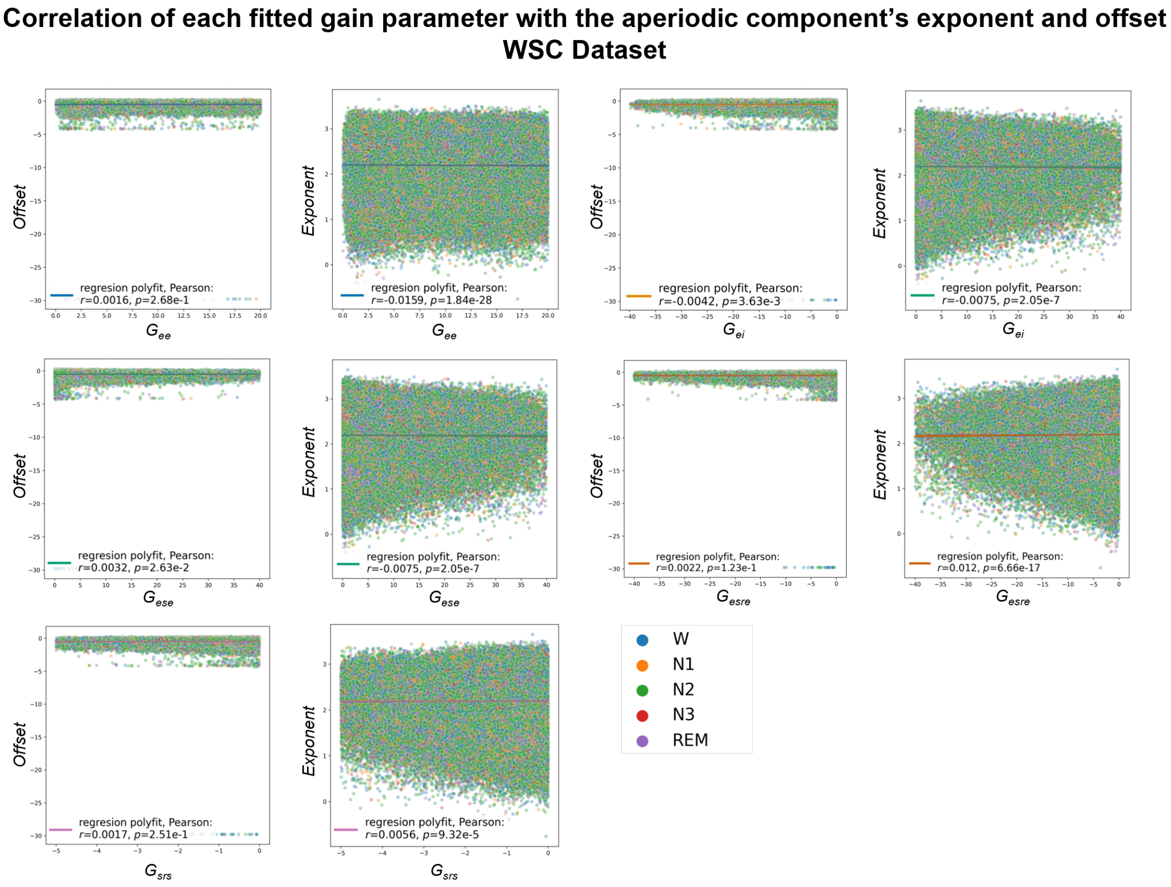


Figure S9: **Associations between gain parameters and $1/f$ components across the WSC dataset.** Points on each plot correspond to individual 30-second epochs throughout the entire dataset, color-coded by sleep stage: Wakefulness (W), NREM Stage 1 (N1), NREM Stage 2 (N2), NREM Stage 3 (N3), and REM sleep (REM). The odd columns from left present correlations with offset values, while the even columns depict correlations with exponent values. Each regression line is accompanied by a Pearson correlation coefficient ($r$) and a $p$-value, indicating the strength and statistical significance of the linear relationship.

##### *The model captures PSDs better than common power spectral band estimation measures*

We also conducted the correlation analyses between the EEG power bands and the fitted parameters from the neurophysiological model. The analysis was conducted as described in the Section 3.2.2 of the paper. The analysis was repeated for all datasets comparing the correlations between the power spectral band powers of the EEG across the commonly-compared frequency bands of delta (0.5 - 4 Hz), theta (4 - 7.5 Hz), alpha (7.5 - 12 Hz), beta (14-25 Hz), and gamma (25 - 45 Hz) with all 9 fitted and calculated model parameters. Correlations were evaluated by using Pearson's $r$-test and False Detection Rate (FDR) correction was done using the Bonferroni method. The $r$-values for these correlations are reported in Figs S10 to S14.

The comparison of these figures further supports the observation in Fig. 4C and 4D that the AUC and FOOOF measurements of the band powers varies greatly and sometimes conflicts due to changes in the broadband power and $1/f$ exponent and peaks all at the same time, where neither of these methods are sufficient to capture the variation in the power spectra alone. In the Dreem & WSC datasets which possess the highest average $1/f$ exponents (per Fig. 2B), this conflict between the AUC and the FOOOF correlations are most evident.

This illustrates the higher significance of broadband and $1/f$ changes as the factors determining the fits when compared to the sharp frequency peaks. This also demonstrates the benefit of using this physiological model fitting method to holistically encapsulate these dynamics that could only partially captured via either of those two methods.

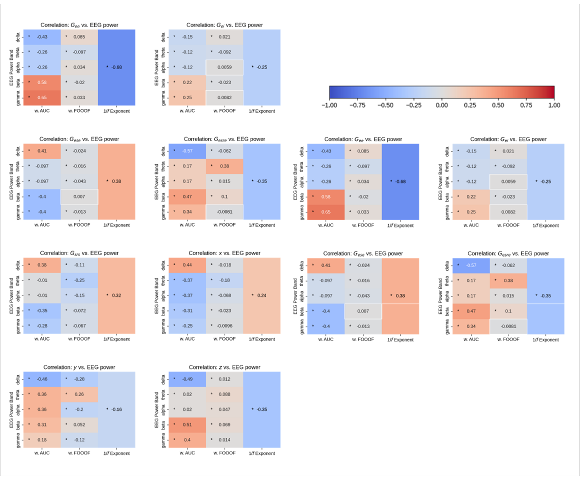

Figure S10: **Correlations between the fitted model parameters and the EEG spectral band powers across the EDF-X dataset** - Band power is calculated via two methods: *AUC*: calculating the area under curve (AUC) by trapezoidal integration of the power spectra, and *FOOOF*: Calculating the height of the above-threshold peaks after separating the scale-free background $1/f$ component from the power spectrum. The significant correlations are highlighted by an asterisk (*) and non-significant correlations are enclosed by a white rectangle.

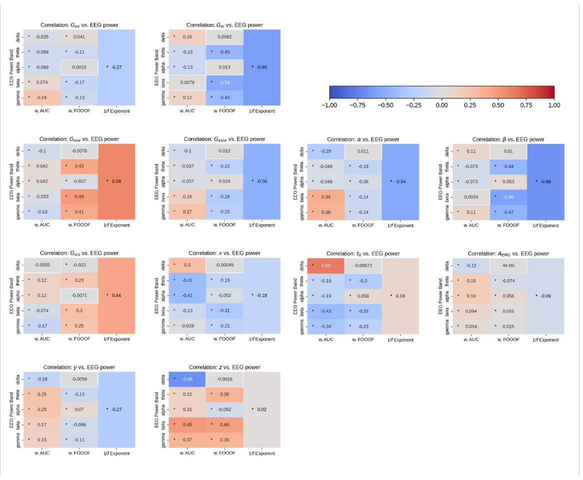

Figure S11: **Correlations between the fitted model parameters and the EEG spectral band powers across the Dreem dataset** - Band power is calculated via two methods: *AUC*: calculating the area under curve (AUC) by trapezoidal integration of the power spectra, and *FOOOF*: Calculating the height of the above-threshold peaks after separating the scale-free background $1/f$ component from the power spectrum. The significant correlations are highlighted by an asterisk (*) and non-significant correlations are enclosed by a white rectangle.

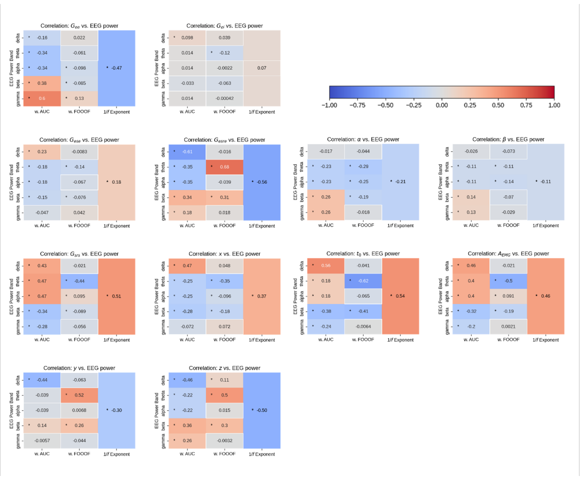

Figure S12: **Correlations between the fitted model parameters and the EEG spectral band powers across the Nap-EEG dataset** - Band power is calculated via two methods: $AUC$: calculating the area under curve (AUC) by trapezoidal integration of the power spectra, and $FOOOF$: Calculating the height of the above-threshold peaks after separating the scale-free background $1/f$ component from the power spectrum. The significant correlations are highlighted by an asterisk (*) and non-significant correlations are enclosed by a white rectangle.

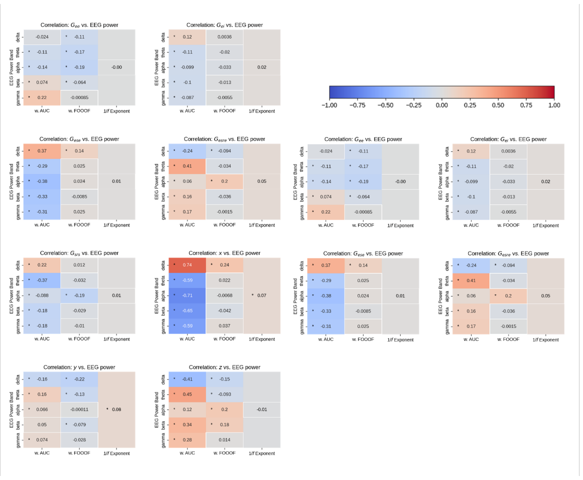

Figure S13: **Correlations between the fitted model parameters and the EEG spectral band powers across the Muse S dataset** - Band power is calculated via two methods: *AUC*: calculating the area under curve (AUC) by trapezoidal integration of the power spectra, and *FOOOF*: Calculating the height of the above-threshold peaks after separating the scale-free background $1/f$ component from the power spectrum. The significant correlations are highlighted by an asterisk (*) and non-significant correlations are enclosed by a white rectangle.

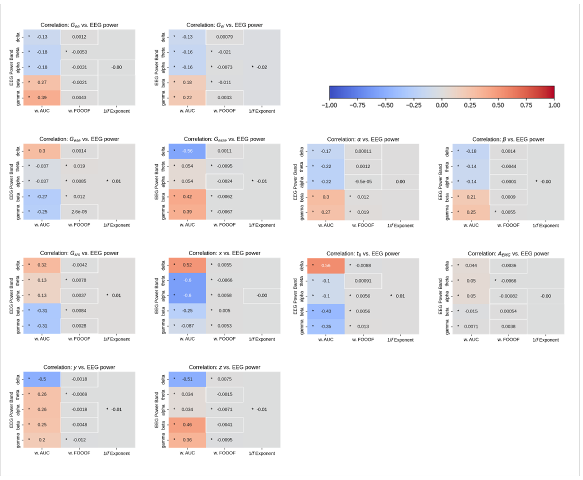

Figure S14: **Correlations between the fitted model parameters and the EEG spectral band powers across the WSC dataset** - Band power is calculated via two methods: $AUC$: calculating the area under curve (AUC) by trapezoidal integration of the power spectra, and $FOOOF$: Calculating the height of the above-threshold peaks after separating the scale-free background $1/f$ component from the power spectrum. The significant correlations are highlighted by an asterisk (*) and non-significant correlations are enclosed by a white rectangle.
